# Cortical long-range dynamics track excitation-inhibition balance across cognitive states and organizational principles

**DOI:** 10.64898/2026.10.30.763069

**Authors:** Matilde Bisi, Asia Ferrari, J. Karolis Degutis, Stefano Tambalo, Angela I. Renton, Michael V. Lombardo, Jorge Jovicich, Dimitri Van De Ville, Francesca Saviola

**Affiliations:** Neuro-X Institute, Ecole Polytechnique Fédérale de Lausanne, Geneva, Switzerland; Faculty of Medicine, University of Geneva, Geneva, Switzerland; Center for Mind/Brain Sciences, University of Trento, Rovereto, Italy; Sant’Anna School of Advanced Studies, Pisa, Italy; Department of Physics, Department of Molecular Biotechnology and Health Sciences, University of Torino, Torino, Italy; Center for Neuroscience and Cognitive Systems, Istituto Italiano di Tecnologia, Rovereto, Italy

## Abstract

A fundamental question in systems neuroscience is how the brain’s excitation-inhibition (E/I) balance shapes the temporal structure of neural activity and whether we can non-invasively recover this relationship from neural scaling properties. We show that MRS-derived E/I balance is expressed in long-range temporal dynamics and constrained by cytoarchitecture. The laminar organisation of inhibitory circuits, spanning agranular to granular cortex, determines how faithfully regional dynamics reflect E/I balance. This neurochemical signature is state-dependent: its relationship with cortical dynamics reverses during task engagement, consistent with traversal of dynamical regimes around a critical point. It extends beyond fMRI to arousal-linked dynamics, indexed by pupil diameter and microsaccades, indicating a global mode of cortical state rather than a regionally isolated property. The imprint of E/I balance on long-range dynamics follows the sensory-association axis at rest, and is reconfigured during task engagement. These findings establish E/I balance as a neurochemical quantity with structured, scalable signatures across the temporal and spatial organization of the human cortex.

## Introduction

The cerebral cortex derives much of its computational flexibility from a single, continuously adjustable quantity: the neurophysiological balance between excitatory and inhibitory neurotransmission (E/I)^1–5^. This homeostatic equilibrium is unlikely to be uniform across the brain. Cortical regions are positioned along a functional hierarchy and differ markedly in cytoarchitectonic properties and cognitive roles; accordingly, E/I balance should be reconfigured across the cortex to support these specialized functions. Whether such large-scale patterning actually exists, however, remains unknown. Perturbations of E/I balance propagate throughout cortical networks and, when maladaptive, are theorized to contribute to a broad spectrum of conditions, ranging from psychiatric and neurological disorders to the cognitive decline associated with aging^6,7^. Establishing whether and how E/I balance is patterned across the cortex and reshaped by ongoing cognitive states would therefore have implications well beyond basic neuroscience, refining mechanistic accounts of these conditions.

A clear methodological roadblock has limited progress on this question: E/I balance has so far been accessible only at discrete cortical sites, leaving its whole-cortex organization and state dependence largely uncharted. Among available techniques, Magnetic Resonance Spectroscopy (MRS)^8–10^ offers the most direct in vivo measure of E/I balance in the human brain by quantifying regional glutamatergic and GABAergic concentrations. Its restriction to a single voxel, however, precludes cortex-wide mapping. In this work, we develop an approach that extends MRS-derived E/I (i.e. Glx/GABA+) characterization beyond single-voxel MRS toward whole-cortex inference. By first clarifying how local MRS-based E/I estimates relate to distributed patterns of ongoing brain activity, we then leverage these relationships to map E/I balance across the cortex and examine its features and modulation by cognitive state.

This strategy is motivated by viewing the brain as a non-linear dynamical system, in which E/I balance acts as a key parameter determining the system’s operating regime. Within neural networks, the E/I ratio tunes cortical dynamics across phase transitions: excessive excitation drives a supercritical state characterized by short-range, unstable activity, whereas excessive inhibition pushes the system into a subcritical, quiescent regime. When finely balanced, activity instead settles near a critical (i.e., *homeostatic*) point, neither quiescent nor unstable, and exhibits structured variability over many scales^11–14^. In this regime, temporal fluctuations become scale-free, obeying a power-law relationship across timescales without a single dominant frequency. The signature of E/I balance should therefore be directly encoded in the aperiodic structure of neural signals.

Diverse approaches, from biophysical modelling^15,16^ to signal-derived statistical measures^14,17^, have sought such non-invasive readouts of E/I in the human brain. One such measurement is the Hurst exponent (H), a measure of long-range temporal dependence that quantifies how persistent or reversible neural fluctuations are over time. Within this dynamical-systems framework, E/I balance (i.e. H) tunes cortical networks along a subcritical-supercritical axis: more inhibition-dominant regimes produce smoother, more persistent activity, whereas more excitation-dominant regimes produce more volatile, rapidly reversing fluctuations. Because H captures precisely this persistence^18–24^, it can serve as a monotonic, inverse surrogate of E/I, linking the aperiodic structure of neural signals to the underlying neurochemical regime^25,26^.

A key question is therefore whether H faithfully tracks MRS-derived E/I balance in humans, and whether this relationship is constrained by cortical architecture and reconfigured by cognitive state. If E/I balance supports region-specific computations, it should vary across the cortex in a manner shaped by cytoarchitecture and functional specialization. Cytoarchitecturally, the cortex is organised along a continuous hierarchy, from agranular regions with thick infragranular layers and diffuse modulatory inhibition to granular regions with a dense layer IV and fast-spiking perisomatic inhibition^27,28^. This gradient determines how precisely inhibition constrains excitatory drive and should therefore constrain the fidelity with which H captures the underlying E/I signal. Functionally, the cortex is further organised along the primary functional gradient, spanning unimodal sensory-motor to transmodal association cortex, a topography linked to the hierarchical organisation of intrinsic neural timescales^27–29^. Behavioural state shifts that alter excitation and inhibition, such as increased arousal^30–34^ during cognitive engagement (operationalizable via cognitive load manipulations and pupillometry measurements), are expected to reconfigure E/I balance, but how such states reshape its cortical expression remains unclear. Whether cytoarchitectural hierarchy and the S-A axis jointly shape the spatial expression of E/I, and whether cognitive demand reconfigures that expression, remains unknown.

Here we use two independent datasets to show that (i) H tracks MRS-derived E/I balance, with the strength of this relationship gated by a region’s position along the cytoarchitectural hierarchy; (ii) the sign of the H-Glx/GABA+ correlation reverses when the brain moves from rest to task engagement, mirroring changes in arousal; and (iii) the cortical topography of H is itself organised by the brain’s S-A gradient, elevated at both the unimodal and transmodal poles at rest, a bipolar structure that selectively changes under cognitive load.

Collectively, our results establish H as a state-sensitive, modality-independent, and structurally constrained readout of cortical E/I regulation. Because H can be estimated non-invasively from whole-brain recordings, it provides a scalable proxy for monitoring disruptions to E/I architecture in conditions marked by inhibitory dysfunction, from neurodevelopmental to neuropsychiatric and age-related disorders.

## Results

### In-vivo mapping of E/I balance proxies

MRS remains the most biologically direct, yet non-invasive, window onto E/I balance in the living human brain. However, its coarse, single-voxel resolution confines Glx/GABA+ (glutamate+glutamine to GABA+) measurements to a single region at a time. This raises an immediate question: can a whole-brain measure substitute MRS, and if so, uniformly across the cortex or only where the underlying cytoarchitecture permits it? We first tackled this question in the discovery WAND dataset^37^, comprising 170 adults scanned with resting-state fMRI and GABA-edited MEGA-PRESS MRS at four voxel locations: left sensorimotor (lSMN), anterior cingulate (ACC), occipital (OCC), and right auditory cortex (rAUD). From resting-state fMRI, we estimated H, indicating how slowly a region’s spontaneous fluctuations decay in autocorrelation, for each subject. Higher H reflect smoother, more persistent dynamics, lower H more rapid, weakly autocorrelated ones. The group-average map revealed that H follows known cytoarchitectural and functional organization: a graded, spatially heterogeneous topography across cortex (Figure 1A) with mean H highest in Default Mode (DMN), Control (CONT), and Dorsal Attention (DAN) networks, and lower in the Limbic (LIM) network. This asymmetry is consistent with LIM’s position at the paralimbic pole of the cortex’s principal microstructural gradient^38^, where diffuse, modulatory inhibition would be expected to support less temporally persistent activity than in the more elaborately laminated association cortex comprising DMN and CONT (Fig. S1). An independent replication cohort of 36 participants (interleaved resting-state/task-based fMRI^10^ with GABA-edited fMRS in dorsolateral prefrontal cortex (DLPFC) during a three load N-back task), showed qualitatively consistent parcel-wise spatial patterns (Figure 1B). To test whether this topography is reproducible across acquisitions and preprocessings, we repeated the analysis in an independent, publicly available dataset, the Human Connectome Project Young Adult dataset^39^ (HCP; n=100 unrelated participants), used as a robustness sample and chosen for its four long resting-state runs per subject, acquired over two sessions. The robustness sample reproduced the same broad cortical topography, with H again lowest in LIM (Figure 1C). Network-averaged radar plots further revealed a consistent pattern across all three datasets (Figure 1D). H differed systematically across the four MRS voxels (Friedman p<0.001, W=0.48), and so did the Glx/GABA+, but in the opposite direction (Friedman p<0.001, W=0.72), region by region: ACC had the lowest H and the highest Glx/GABA+, whereas lSMN the highest H and the lowest Glx/GABA+ (Fig. 1E). This inverse, region-by-region correspondence indicates resting-state H recovers regional spectroscopic information rather than diffuse, whole-brain signal (Fig. S2). This pattern held in the replication sample, despite a different cohort, scanner, sequence and regions: resting DLPFC H and Glx/GABA+ fell within the discovery voxels’ range (Fig. 1F). Parcel-wise H topographies converged: all cross-cohort matrix correlations were significant (discovery-replication: r = 0.30, p_spin_ = 0.001; discovery-robustness: r = 0.53, p_spin_ = 0.001; replication-robustness: r = 0.35, p_spin_ = 0.001; Fig. 1G), showing reproducible fine-grained H covariance across cohorts and protocols. Test-retest reliability across sessions is reported in the Supplementary Results.

**Figure 1.**
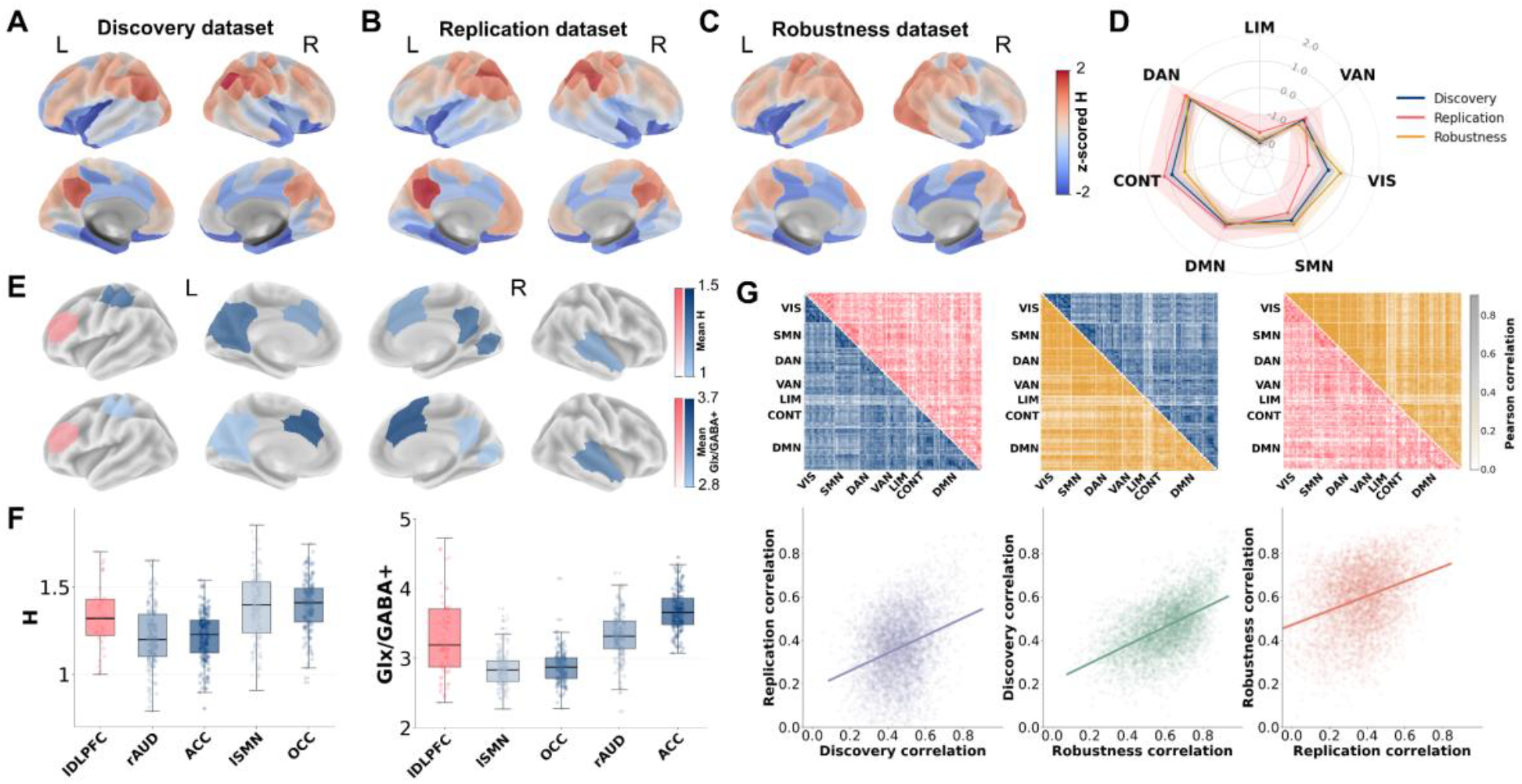
Whole-cortex correspondence between resting-state BOLD fMRI Hurst exponent (H) and MRS excitation-inhibition balance is constrained by cortical cytoarchitecture and replicated across independent cohorts. **A**. Z-scored mean parcel-wise Hurst exponent maps at rest in the discovery dataset. Darker colours indicate higher H and lighter colours indicate lower, more antipersistent H. **B**. Z-scored mean parcel-wise Hurst exponent maps at rest in the replication dataset. **C**. Z-scored mean parcel-wise Hurst exponent maps at rest in the robustness dataset. **D**. Z-scored network-averaged mean H displayed as radar plots for discovery (blue shading), replication (pink shading) and robustness (yellow shading) datasets. Shaded areas in the radar plot indicate 95% CI. **E**. Mean H and Glx/GABA+ ratio maps for the four MRS voxels in the discovery dataset (blue shading) and the DLPFC voxel in the replication dataset (pink shading). **F**. Boxplots of H and Glx/GABA+ across MRS voxels in the discovery and replication datasets. **G**. Parcel-by-parcel Pearson correlation matrices of H across subjects in the discovery (blue), replication (pink), and robustness (yellow) samples. Each split matrix displays the correlation structure of a pair of datasets in its upper and lower triangles, with the corresponding scatter plot and regression line below. Abbreviations: VIS, Visual; SMN, Somatomotor; DAN, Dorsal Attention; VAN, Salience/Ventral Attention; LIM, Limbic; CONT, Control; DMN, Default Mode Network; lDLPFC, left Dorsolateral Prefrontal Cortex; ACC, Anterior Cingulate; OCC, Occipital Cingulate Cortex; rAUD, right Auditory; lSMN, Somatomotor left.

Together, these findings indicate that whole-brain correspondence between H and Glx/GABA+ identified in the discovery dataset is not idiosyncratic to a single cohort or acquisition protocol.

### State-dependent association between BOLD scale-free dynamics and Glx/GABA+

Resting-state correspondence between BOLD-derived H and Glx/GABA+ does not, by itself, establish whether this coupling is a fixed biological signature or also tracks cognitive engagement shifts. Prior work (Bertelsen et al.,) suggests that the relationship between H and synaptic E/I conductance ratio may reverse as a function of input excitability to local neural networks. The distinction between rest versus task would be a manipulation heavily altering input excitability. Based on these ideas, the relationship between H and Glx/GABA+ should change across rest versus task conditions. We tested this within the same MRS voxel across four sessions of working-memory N-back task (i.e. rest and three increasing loads (0-, 1-, 2-Back)) which place progressively greater demand on holding, updating and inhibiting no-longer relevant items in working memory (Figure 2A-B). Mean Glx/GABA+ was comparable across sessions, with overlapping distributions between rest and N-back conditions (Figure 2C): Glx/GABA+ itself was not state-dependent, yet the Glx/GABA+-H relationship differed across cognitive states. A significant Glx/GABA+ × Session interaction reflected a reversal: at rest, higher Glx/GABA+ was associated with lower H (β=−0.16, pFDR<0.001), consistent with suppression of temporal persistence, whereas under any working-memory load the relationship flipped to positive and grew stronger with load (0-Back: β=+0.13, pFDR=0.002; 1-Back: β=+0.16, pFDR<0.001; 2-Back: β=+0.22, pFDR<0.001), suggesting increases in signal persistence. This sign reversal converges with in-silico modeling of coupled excitatory-inhibitory networks26 in which the relationship between an aperiodic, H-like signal component and the synaptic E/I conductance ratio itself flips sign as a function of input excitability. Increasing cognitive load plausibly raises input excitability to prefrontal circuitry is a comparable way, shifting DLPFC from a rest-like asynchronous irregular firing regime towards a more synchronous irregular firing regime as working-memory demands increase. Decomposing Glx/GABA+ into GABA+ and Glx did not reproduce this interaction for either metabolite, suggesting the composite MRS-derived E/I index captures a state-dependent relationship not evident in individual metabolites (Supplementary Fig. S3).

**Figure 2.**
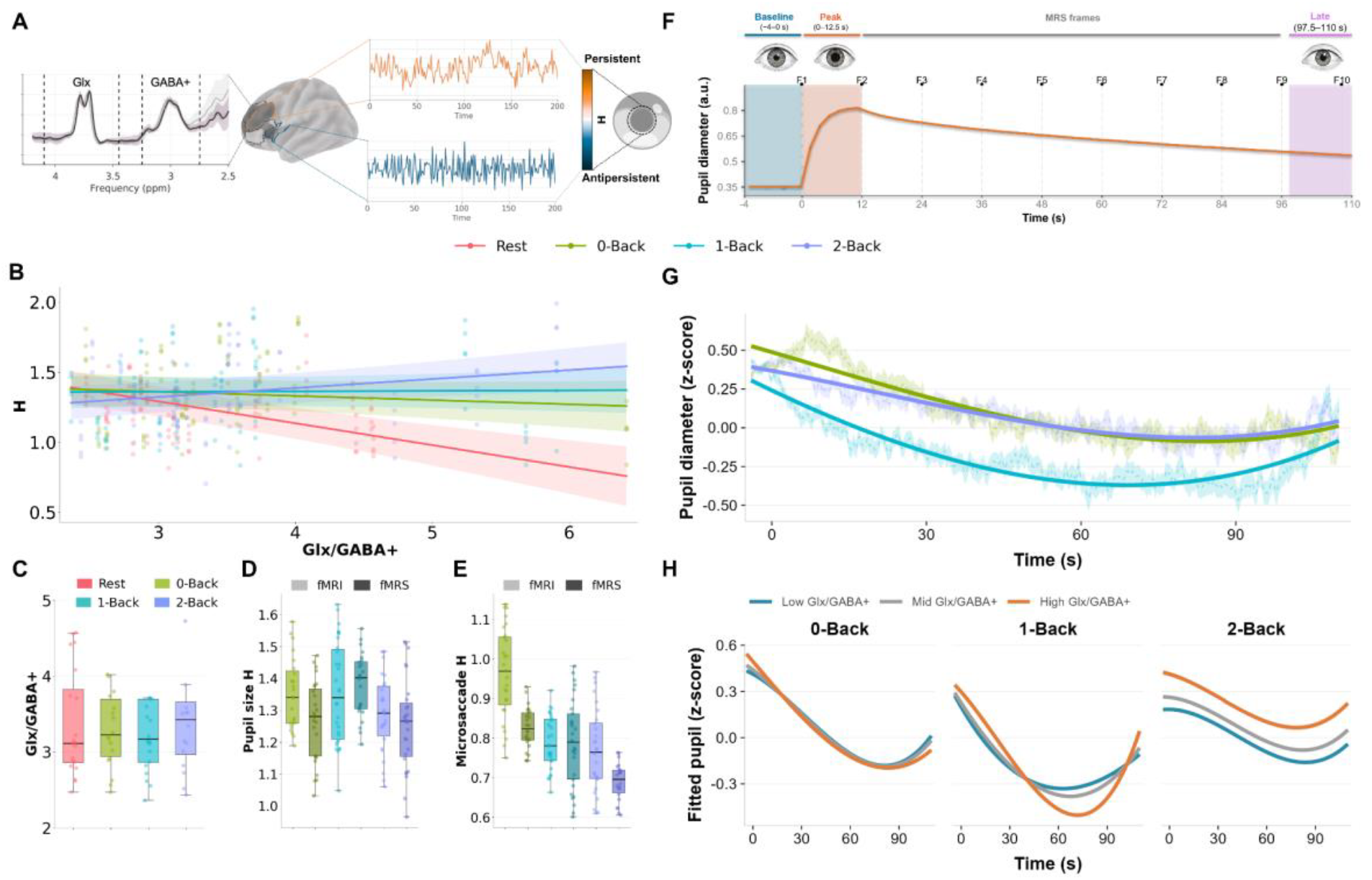
Excitation-inhibition balance as a state-dependent predictor of cortical Hurst exponent and pupil dynamics. **A**. Schematic overview of the methodology. fMRS was used to estimate Glx/GABA+ ratio in a left dorsolateral prefrontal cortex voxel, whereas Hurst exponent was computed from parcel-wise BOLD signals and pupillometry data. **B**. Relationship between BOLD-derived Hurst, averaged across the parcels comprising the left DLPFC MRS voxel, exponent and Glx/GABA+ across sessions. Lines represent fixed effects from a linear mixed-effects model of the form H ~ Glx/GABA+ × Session + (1 | subject + parcel). **C**. Boxplots of Glx/GABA+ across sessions. **D**. Boxplots of pupil size-derived H for all sessions. **E**. Boxplots of microsaccade-derived H for all sessions. **F**. Theoretical polynomial curve fitted to the group-mean pupil diameter trajectory (arbitrary units), illustrating the canonical waveform used to parameterize growth curve analysis (GCA) models. The pre-task baseline window (−4 to 0 s) and late epoch (97.5-110 s) are shaded in grey. Red dashed lines mark ten equidistant time frames (F1-F10), each corresponding to an MRS acquisition frame within the block epoch. **G**. Mean observed (dashed, ±1 SE shaded) and GCA-1 model-fitted (solid) pupil diameter trajectories across the three N-back conditions. **H**. Fitted pupil trajectories from the three-way interaction model (GCA-3: polynomial time × Glx/GABA+ × Task), displayed separately for each task condition and stratified by static Glx/GABA+ tertile.

In sum, we found that H’s relationship to Glx/GABA+ is state-dependent rather than fixed: the monotonic, inverse coupling evident at rest reverses under cognitive engagement. This establishes H not as a static marker of regional Glx/GABA+ but as a dynamic readout of its reconfiguration under cognitive demand.

### Arousal-dependent coupling between H and Glx/GABA+

If the state-dependent H–Glx/GABA+ coupling reflects a shift in global brain state, related signatures may extend to peripheral readouts of arousal. Two such signals are pupil diameter and microsaccade rate; both well established to reflect cognitive load and arousal^37–39^. We used the eye-tracking data available in the replication dataset to ask whether temporal structure of arousal fluctuations (i.e. H derived from pupil size and microsaccade rate time series) also change with cognitive load.

Both signals habituated across the six fMRI/fMRS runs, as expected from simple vigilance decrement (Figure 2D-E), but this decline was partially offset under moderate load: pupil-derived H showed an attenuated decline under 1-Back (β=+0.11, p=0.025), while microsaccade-derived H was buffered under both 1-Back (β=+0.13, p<0.001) and 2-Back (β=+0.07, p=0.02), indicating that microsaccadic persistence resisted habituation across a wider range of cognitive load than pupillary H (see Supplementary Results). This suggested that the two peripheral channels are not redundant readouts of the same arousal signal.

Engagement reorganized the pupil’s moment-to-moment behaviour within each block. A growth curve analysis (GCA) of the pupil waveform showed the expected signature of task-driven arousal followed by habituation (i.e. an early peak, sustained decline and a partial late recovery). This shape was modulated by load (Figure 2G): 1-Back and 2-Back both attenuated the initial decline relative to 0-Back, while 1-Back additionally deepened and delayed the recovery trough. Glx/GABA+ balance further shaped this trajectory in a load-dependent manner (Figure 2H): under 1-Back it predicted the shape of the pupil response itself, whereas under 2-Back it predicted a sustained elevation in overall pupil diameter instead. This indicates that Glx/GABA+’s expression in arousal dynamics shifts from temporal structure to tonic level as cognitive demand increases (see Supplementary Results). However, applying the same Glx/GABA+-by-session framework to the scalar H summaries of pupil and microsaccade signals yielded no significant effects, neither as the composite ratio nor when decomposed into GABA+ and Glx (Fig. S4), likely reflecting the reduced sensitivity of a single scalar summary, relative to the time-resolved GCA above.

Together, we found that Glx/GABA+’s influence on arousal is not confined to the cortex: peripheral signals under prefrontal control carry a comparable, load-dependent signature, expressed as a buffered habituation and a reorganized pupil trajectory that shifts from temporal structure to tonic elevation under highest load.

### E/I proxies are sensitive to cognitive load shifts

H’s resting-state topography tracks MRS-derived E/I changes regionally; we next tested whether the whole-brain organization itself reorganizes under cognitive engagement. We estimated the H parcel-wise in each condition (Figure 3A). Whole-cortex mean H remained broadly stable across conditions (rest =1.26±0.15; 0-Back=1.30±0.10; 1-Back=1.31±0.11; 2-Back=1.24±0.11), with no monotonic modulation across increasing N-back load, indicating that rest-to-task transition, rather than graded demand, primarily drives global cortical reconfiguration. This stability, however, concealed marked spatial heterogeneity (detailed voxel-wise and parcel-wise statistics in Supplementary Fig. S5, S6 and Table S2): reconfiguration was carried by specific networks rather than uniformly distributed across cortex (Figure 3B). A significant Network×Session interaction (F_(18,594)_=13.15, p_GG-cor_r<0.001, partial η^2^=0.28) confirmed that task reorganizes cortical persistence unevenly across networks. Parcel-wise ANOVAs identified significant session effects in 49% of parcels (p_FDR_<0.05), with the strongest effects (partial η^2^>0.3) concentrated in LIM and DMN (Figure 3C). LIM (encompassing both orbitofrontal and temporal poles) carried the clearest signature: the lowest H of any network at rest (0.98±0.22) and the largest task-related increase. CONT (including dorsolateral prefrontal and posterior parietal cortex) and DMN (comprehending medial prefrontal cortex, precuneus and angular gyrus) maintained the highest H throughout (mean across sessions: DMN=1.30±0.25; CONT=1.36±0.25), consistent with their role as higher-order integration hubs. On the other hand, unimodal sensory and attentional networks shifted only modestly. As at the whole-cortex level, session ordering was non-monotonic, with 1-Back and 0-Back consistently exceeding 2-Back, indicating that moderate rather than maximal load drives the greatest cortical persistence. Detailed network-level analyses, including the Network × Session repeated-measures ANOVA and pairwise comparisons, are reported in the Supplementary Results (Tables S3-S8; Figure S7).

**Figure 3.**
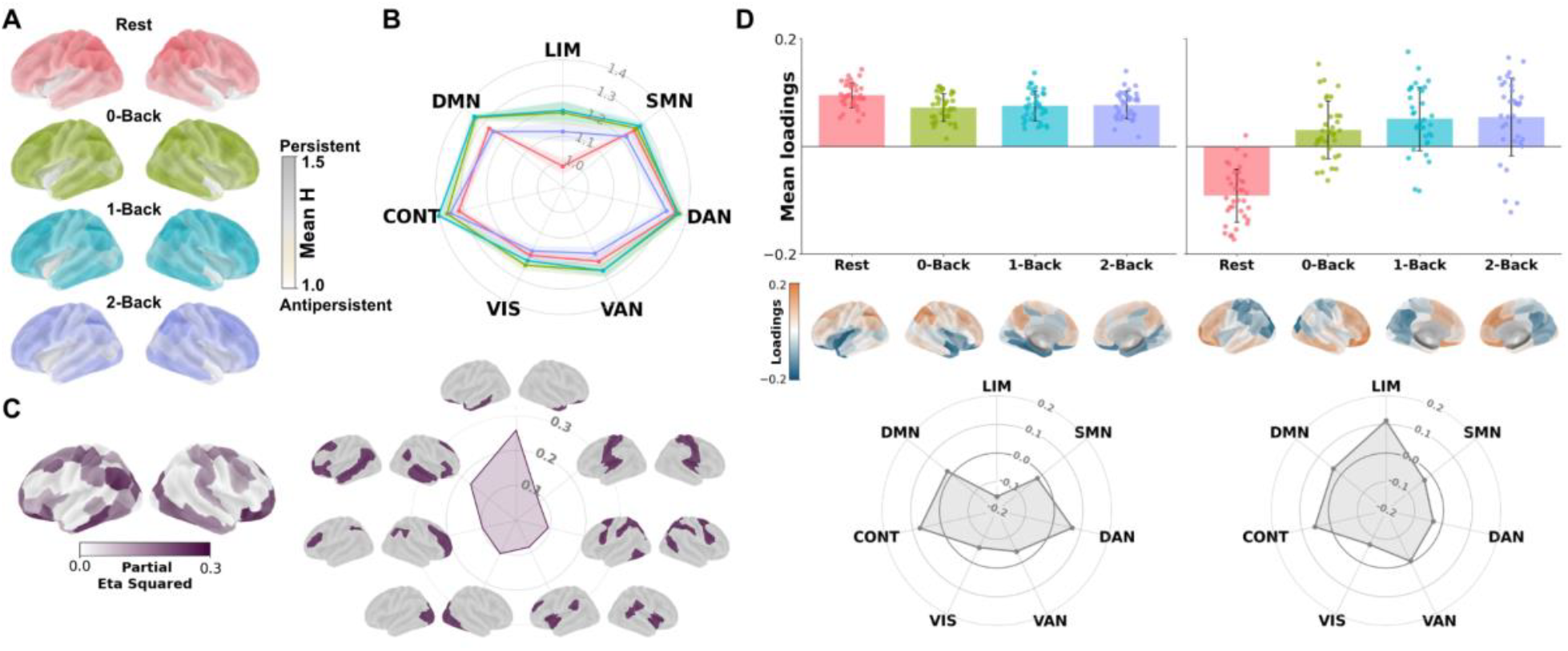
Cortical long-range dynamics follow the brain’s intrinsic functional gradient and reorganize under task. **A**. Mean parcel-wise Hurst exponent maps across Rest, 0-Back, 1-Back, and 2-Back. Darker colours indicate higher H and lighter colours indicate lower, more antipersistent H. **B**. Network-averaged mean H displayed as radar plots for the seven functional networks across the four sessions. Shaded areas in the radar plot indicate 95% CI. **C**. Parcel-wise partial eta squared maps from the repeated-measures ANOVA, together with the corresponding network-level summary of effect sizes. **D**. Singular value decomposition of the parcel × (subject·session) H matrix, showing the mean session loadings of the first two dominant components, their associated spatial loading maps and the mean network-wise loadings as radar plots. Abbreviations: VIS, Visual; SMN, Somatomotor; DAN, Dorsal Attention; VAN, Salience/Ventral Attention; LIM, Limbic; CONT, Control; DMN, Default Mode Network.

To complement these univariate, network-defined analysis with a data-driven multivariate perspective, we applied singular value decomposition (SVD) to the mean-centered H values, arranged with cortical parcels along one dimension and each subject’s individual task sessions along the other. While the network-based analysis identified univariate differences in H at predefined regions, it could not reveal whether these regional differences reflected a spatial pattern that was coherently shared across the cortex. SVD allowed us to extract common spatial patterns of H variability across parcels and task conditions, and to identify which of these patterns were consistently expressed across subjects. This decomposition yielded two dominant components explaining 40% and 9% (p<0.001) of total variance (Figure 3D).

The first captured an intrinsic spatial mode, strongest at rest and modestly attenuated during task, with LIM at one pole (lower H) and CONT/DAN at the other (higher, more persistent H). The second showed a clearer rest-to-task gradient (i.e. state-dependent), rising monotonically from rest through increasing load, with an opposing spatial pattern: H rose during task in sensory and attention regions (VIS, SMN, DAN) and fell in DMN and CONT. This decomposition confirmed, independent of a priori network boundaries, that the network-level pattern reflected two dissociable sources of variance: (i) a stable spatial hierarchy, and (ii) an orthogonal state-dependent gradient, not recoverable from session-wise contrasts alone. The stable component aligns with the primary functional gradient (|r|=0.28, p_spin_=0.03, 10000 rotations; Supplementary Fig. S8), seen as the principal hierarchy organizing cortical, microstructure, connectivity and function in the brain, places H’s trait topography on the brain’s fundamental organizational cortical principle.

In sum, we found that cognitive engagement reorganizes cortical persistence selectively rather than globally: whole-cortex H remains stable, but LIM and DMN carry the clearest task-related shifts, decomposing into two dissociable spatial modes. On one hand, an intrinsic hierarchy anchored to the primary functional gradient, and on the other an orthogonal, state-dependent gradient emerging only under task conditions.

### The coupling between cortical dynamics and Glx/GABA+ is organised along structural and functional hierarchies

The preceding sections trace where H can track Glx/GABA+ and how that tracking reorganizes with task; in what follows we investigate what mechanism underlies this correspondence. Specifically, we asked whether the coupling between MRS-derived E/I and cortical long-range dynamics is uniform across the cortex, or whether the sign and strength of the Glx/GABA+-H relationship is itself dependent on the underlying anatomy. Cortical regions differ systematically in laminar organisation, which determines how local inhibitory circuits are organised and how precisely they shape excitatory drive: a plausible structural mechanism constraining where an Glx/GABA+-H relationship should exist at all. To test this, we characterised the constituent parcels of each MRS voxel in the discovery dataset along two independent cytoarchitectural frameworks using linear mixed-effects models^43,44^.

The Glx/GABA+–H relationship was not uniform across cytoarchitecture, but depended on where a region sat along the cortical hierarchy. Using the continuous BigBrain45 microstructural gradient (Figure 4A, third-order polynomial as the superior fit), the model showed significant effects of Glx/GABA+ (β1 = −0.2, p < 0.001) and of the gradient itself (β1 = −0.6, p < 0.001), together with a significant Glx/GABA+×Gradient interaction (β2 = −2.0, p = 0.015).

**Figure 4.**
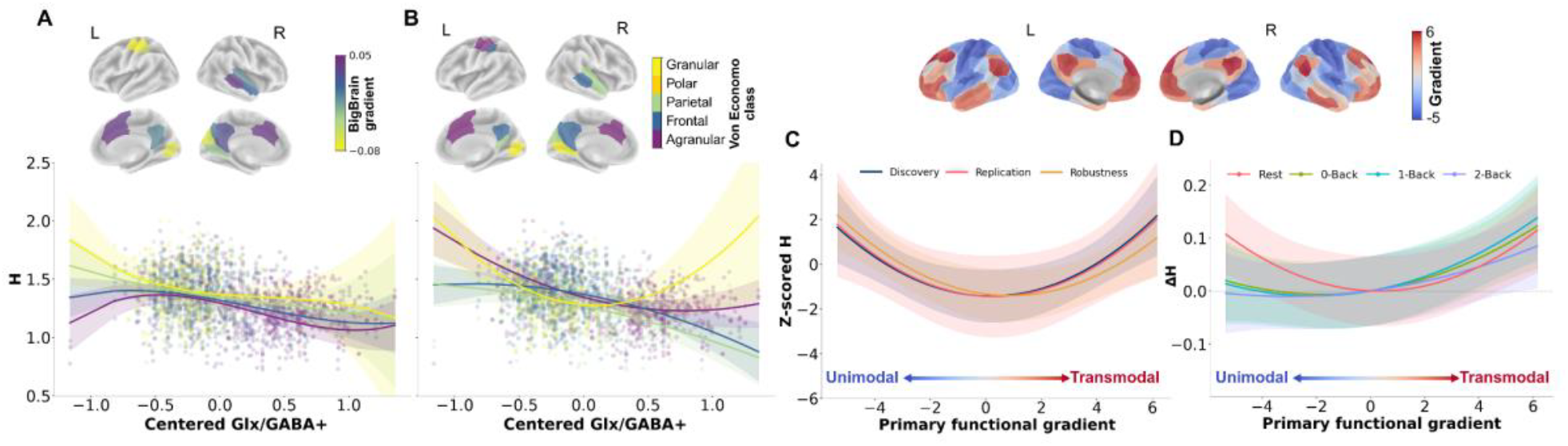
Glx/GABA+ and H reflect macroscale cortical organization. **A**. Fitted Glx/GABA+ from the polynomial models with von Economo cytoarchitectural information across the MRS voxels of the discovery dataset. **B**. Fitted Glx/GABA+ from the polynomial models with BigBrain gradient across the voxels. **C**. Z-scored predicted values of H as a function of the principal gradient of cortical organization (Margulies et al., 2016) in the discovery, replication and robustness samples fitted from quadratic mixed-effects models (shaded band, 95% CI). **D**. Mean-centered predicted values of H for different sessions as a function of the principal functional gradient in the replication sample fitted from a quadratic mixed-effects model with interaction between gradient and session.

To characterize this interaction, we then examined the local Glx/GABA+–H slope at the two ends of the cytoarchitectural gradient, evaluated at mean-centered Glx/GABA+ = 0. The slope was steep and significant in associative cortex (slope = −0.25, pFDR = 0.003), but weaker and non-significant at primary sensory anchors (slope = −0.10, pFDR > 0.05).

The same pattern held using discrete von Economo-Koskinas44 cytoarchitectonic classes (Figure 4B, second-order polynomial as superior fit), with a significant Glx/GABA+ × Cytoarchitectonic-class interaction (β2=0.4, p<0.001) alongside main effect of Glx/GABA+ (β1=−0.2, p<0.001; β2=0.1, p=0.004) and class (β1=−0.07, p<0.001). Together, these results show that Glx/GABA+-H coupling is not uniform but strongest in associative regions and negligible at primary sensory anchors, indicating a hierarchically graded cytoarchitectural constraint (Table S9 and Fig. S11).

Beyond recovering region-specific E/I information constrained by cytoarchitecture, H’s spatial organization also recapitulates the intrinsic functional hierarchies, the so-called primary functional gradient, that organize large-scale cortical activity, indicating this is a general property of its topography rather than one specific to this task. Relating resting-state H to the principal gradient of macroscale cortical organization^29^, H traced a U-shaped function of the gradient in both cohorts, lowest near the gradient’s center, rising toward unimodal and transmodal poles (quadratic fit superior to linear in both discovery: χ^2^(1)=9.88, p=0.002 and replication: χ^2^(1)=4.24, p=0.039, Figure 4C) indicating that cortical long-range dynamics are organized along the same principal axis governing large-scale functional connectivity. This U-shaped organization was independently reproduced in the HCP dataset (χ^2^(1)=10.00, p<0.01). The relationship between H and functional hierarchy further reorganizes under cognitive engagement, extending the state-dependence seen throughout this dataset. The U-shape flattened into a more linear form as load increased (quadratic term significant at rest, 0- and 1-Back but not 2-Back, linear term significant at every load except rest; Figure 4D). This change was driven by one end of the gradient: at rest, H decreased with the primary functional gradient at the unimodal pole (β=−0.029, p=0.003) and increased at the transmodal pole (β=0.025, p=0.004), a clear asymmetry (p=0.002) disappearing under the highest load. Task engagement overrides sensory cortex’s intrinsic, hierarchy-linked temporal dynamics while leaving association cortex’s untouched. Both linear and quadratic gradient components differed significantly from rest across all task loads, confirming load-dependent reconfiguration of the H-gradient relationship (full statistics in Supplementary Results).

In sum, we found that the coupling between cortical dynamics and E/I is organized along both structural and functional hierarchies: constrained by cytoarchitecture, it is strongest in the associative cortex and negligible at primary sensory anchors, following a U-shaped trajectory along the S-A that flattens under cognitive load. Where H tracks MRS-derived E/I, and how faithfully, is therefore set by the cortex’s underlying anatomical and functional organization rather than uniform across the brain.

## Discussion

The fundamental goal of this study was manifold, but centered around the core concept of building a neurovascular readout able to efficiently capture cortical E/I. First, we tested whether the Hurst exponent (H), could capture spectroscopy-based estimates of cortical E/I balance in two independent cohorts, and showed that neurovascular dynamics carry a robust and reproducible signature of cortical E/I balance generalizable in datasets without MRS. Second, we investigated whether this signature follows common organisational principles, shaped by cortical cytoarchitecture along the functional hierarchy, across resting-state and cognitive conditions. Furthermore, we found that the strength of the coupling between H and MRS-derived E/I depends on cytoarchitecture, and that its direction reverses between rest and cognitive engagement. By linking neurovascular dynamics to neurochemical balance^10^, this work connects local microcircuit physiology to large-scale cortical dynamics and cognition. This positions us to learn how the organising principles of the human brain, from laminar architecture to brain state, shape the expression of E/I balance in cortical dynamics, and to advance our understanding of how E/I balance supports healthy brain function and how its disruption may contribute to neuropsychiatric disorders^46,47^.

In terms of mechanistic insights, these findings empirically exemplify, across modalities, cohorts, and organizational axes, a framework that provides a neuroscientific operationalization of H. In this framework, H is a proxy for the excitability of a network, and its relationship to E/I balance is conditional on the input regime in which the circuit operates^48,49^. Seminal work has established a relationship between E/I and H both at rest and in clinical populations^47–50^, largely inferred from the link between the aperiodic, scale-free component of neural signals and excitatory-inhibitory coupling^3,54^. Here, we show that the sign of this relationship reverses across cognitive states in a young healthy population. At rest, higher Glx/GABA+ predicts lower H: individuals with stronger excitatory drive relative to inhibition show reduced temporal persistence. Under cognitive load, the relationship inverts, and higher Glx/GABA+ predicts greater H, consistent with an excitation-driven extension of signal memory. Mean Glx/GABA+ did not differ between rest and task, so what changed was not Glx/GABA+ itself but the regime in which it is expressed. This is what computational modelling^26^ of H against synaptic E/I conductance ratio (g, where g = g_EE_/g_IE_; g_EE_ are synaptic conductances of E neurons on other E neurons; g_IE_ are the synaptic conductances of I neurons on E neurons predicts across input regimes. In that model, H is, first of all, a proxy for the overall excitability of the network, defined as its firing rate: as firing rates rise, H falls. The balance between excitatory and inhibitory conductances then nudges H around this baseline, and the direction of the nudge depends on how strongly the network is driven. At low levels of input excitability, networks are locked into an asynchronous-irregular (AI) firing regime. Under this AI regime, H and g are negatively related. However, as input excitability rises, local networks reconfigure into a synchronous-irregular (SI) firing regime. Under SI regimes, the H-g relationship turns positive. In our work, the transition from rest to task is expected to shift input excitability substantially, particularly in a region such as the DLPFC that is driven hard by working-memory demands^55^, and so accounts for the reversal we observe. Rest and task therefore do not merely differ in the arousal of the participant, but occupy distinct dynamical regimes, with the sign of the H-Glx/GABA+ relationship tracking their separation.

This has practical implications for how H should be interpreted. At rest, higher Glx/GABA+ was associated with lower H in two independent cohorts, and the underlying topography of H was reproduced in a third, independent dataset (HCP) acquired with a different scanner and preprocessing, which offers a working assumption for resting-state data. Under task conditions, or whenever groups or sessions plausibly differ in input drive, the sign cannot be taken for granted, and H is best read alongside an independent index of the state of the system. Pupil size and microsaccade rate are natural candidates: both are established markers of arousal and cognitive load, both are regulated through circuits under prefrontal control, and both showed load-dependent H in our data, as discussed below. The account does not tie the reversal to working-memory load: any condition that shifts input excitability could in principle move a region between regimes, and with it the sign. Whether the reversal extends beyond cognitive load remains to be tested.

The spatial organisation of the coupling follows the same logic. The relationship between Glx/GABA+ and H was not uniform across the cortex: it was steepest in association cortex and negligible at primary sensory anchors, whether cytoarchitecture was described along the continuous BigBrain gradient or by discrete cytoarchitectonic classes. Cortical regions differ systematically in laminar organisation and in the circuitry through which inhibition shapes excitatory drive, and the intrinsic timescales of activity lengthen along the same sensory-to-association hierarchy^27,31^. Where activity is dominated by feedforward input, as in primary sensory cortex, the temporal structure of the signal is plausibly set largely by that input, leaving little room for local E/I balance to express itself; in association cortex, where recurrent circuitry sustains activity over longer periods, E/I balance becomes a stronger determinant of long-range temporal dependence. This reading is consistent with the input-excitability account above: in both cases, how much of the Glx/GABA+ balance is expressed in H depends on how strongly a circuit is driven from outside relative to how much it is shaped from within. It also fits our observation that task engagement overrides the hierarchy-linked dynamics of the sensory cortex while leaving those of the association cortex largely untouched.

H is not, however, a restatement of the functional gradient which is largely considered the main principle of functional organization. Its resting-state topography traced a U-shaped function of the primary functional gradient in both cohorts, lowest near the centre of the gradient and rising towards the unimodal and the transmodal pole, whereas a measure that simply duplicated the gradient would be expected to vary monotonically along it; the stable spatial component of H aligned with the axis only moderately (|r|=0.28). Under cognitive load the U-shape flattened, and this change was carried by the unimodal end of the axis: the asymmetry between the two arms of the U present at rest disappeared at the highest load. That task engagement alters the dynamics of sensory cortex while leaving those of association cortex largely untouched agrees with reports of a task-related reduction of gradient slope confined to unimodal cortex, and with the input-driven account above, since sensory cortex is the part of the hierarchy whose temporal structure should be most readily overridden by task-related input.

Moreover, the sensitivity of H to cognitive state is not specific to the BOLD signal. H of slow pupil fluctuations and of microsaccade rate also responded to load: both signals habituated over the session, and moderate load partly offset this decline, in pupil-derived H under 1-Back and in microsaccade-derived H under both 1- and 2-Back, so that the two channels are not redundant readouts of a single arousal signal. This is plausible in light of the circuits involved. Pupil size indexes the pupil-linked arousal system, including the locus coeruleus-noradrenaline system, which is under top-down control from the prefrontal cortex^53–56^, whereas microsaccades arise from the rostral superior colliculus, whose activity is linked to the locus coeruleus and modulated by attention and task difficulty^38,57–60^. Both effectors are therefore regulated by the prefrontal systems in which we measured Glx/GABA+ balance, which offers a route through which prefrontal state can be expressed in peripheral signals. In keeping with this, Glx/GABA+ was related to the time-resolved pupil response in a load-dependent manner, shaping its trajectory under 1-Back and its tonic level under 2-Back, whereas no Glx/GABA+ effect was found on the scalar summaries of H, probably because a single value per session is less sensitive than the time-resolved analysis. The non-monotonic effect of load, with partial recovery of H under moderate but not under the highest load^13,64,65^, is not what a simple scaling of H with arousal would predict, which points to load-specific effects beyond arousal.

## Limitations

Several limitations should be considered. Glx/GABA+ balance was estimated by MRS in a small number of locations, four voxels in the discovery cohort and a single one, in the DLPFC, in the replication cohort, so that the gradients along which the coupling varies rest on a limited sampling of the cortex. The design is correlational and, for Glx/GABA+, cross-sectional between participants: mean Glx/GABA+ did not differ between sessions, so within-person changes in Glx/GABA+, such as those produced by pharmacological challenge, remain to be tested. The Glx/GABA+ ratio is a composite of metabolite pools that does not distinguish synaptic from metabolic or extracellular contributions, and decomposing it into its components did not reproduce the state-dependent interaction. H is derived from BOLD, a haemodynamic signal, and the task-related reversal could in part reflect changes in neurovascular coupling or arousal that our design cannot separate from changes in neural input, although the match with the input-regime predictions of the computational model offers independent support for a neuronal origin. Finally, H estimates depend on acquisition, preprocessing and time-series length: we therefore interpret relative differences across regions, states and individuals, and not absolute values, and our cohorts comprise young healthy adults, so that generalisation to clinical populations remains to be shown.

## Conclusions

In summary, the temporal structure of cortical activity carries a readout of E/I balance whose strength is shaped by cortical cytoarchitecture and functional hierarchy, and whose direction is set by brain state. Calibrated against spectroscopic estimates of E/I balance in two independent cohorts, H thus becomes operational: a measure with a long-established theoretical meaning that can now be read in terms of network excitability and, conditionally, E/I balance, provided one knows where in the cortex and in what state it is measured. Because resting-state and task fMRI are widely available, this opens cortex-wide questions that single-voxel spectroscopy cannot address: how E/I balance is organised across the cortex, how its expression changes with cognitive state, and how it differs across populations, including in datasets acquired without MRS. In clinical cohorts, the state dependence defines the conditions of use, with resting-state data as the best-characterised starting point.

## Materials and Methods

### Discovery sample: the Welsh Advanced Neuroimaging Database (WAND)

#### Participants (WAND)

To assess the generalizability of the association between H and Glx/GABA+ at rest in an independent cohort, we replicated the discovery analysis in the Welsh Advanced Neuroimaging Database (WAND), a publicly available multimodal dataset of 170 healthy adult volunteers (aged 18-63 years, median 25) recruited at the Cardiff University Brain Research Imaging Centre (CUBRIC). Ethical approval was granted by the Cardiff University School of Psychology Research Ethics Committee (EC.18.08.14.5332RA3) and all participants provided informed written consent. The dataset is organized in Brain Imaging Data Structure (BIDS) format and distributed via the GIN repository (https://gin.g-node.org/CUBRIC/WAND).

#### MR acquisition and processing (WAND)

All data used here were acquired on a 3 T Magnetom Prisma (Siemens Healthcare) with a 32-channel receive-only head coil. From ses-03 we used the T1-weighted MPRAGE (TR/TE = 2250/3.06 ms, TI = 850 ms, flip angle = 9°, 1 mm isotropic, FOV 256 × 288 × 176 mm^3^, 2-fold in-plane undersampling, A≫P phase-encoding) and resting-state multiband BOLD-EPI (TR/TE = 2000/30 ms, flip angle = 70°, 2 × 2 × 2 mm^3^ voxel, FOV 192 × 192 × 160 mm^3^, multiband factor 4, 2-fold in-plane undersampling, A≫P; 10 min 38 s with fixation on a central white cross). From ses-05 we used the GABA-edited MEGA-PRESS spectra (TR/TE = 2000/68 ms, 160 excitations per spectrum, spectral width 2000 Hz, 2048 spectral points, VAPOR water suppression; editing pulses at 1.9 and 7.5 ppm, 100 Hz bandwidth). MRS was acquired from four single-voxel locations per participant: left sensorimotor (lSMN) (30 × 30 × 30 mm^3^), anterior cingulate (ACC) (40 × 30 × 25 mm^3^), occipital (OCC) (30 × 30 × 30 mm^3^) and right auditory cortex (rAUD) (45 × 30 × 20 mm^3^) each accompanied by a matched non-water-suppressed reference acquisition. The TR/TE, editing scheme and voxel volumes closely matched the discovery acquisition, enabling direct methodological replication. MRS preprocessing used Gannet66 3.2.0: spectra were loaded with frequency-and-phase correction (robust spectral registration), ON/OFF subtraction, weighted averaging, and residual-water removal via HSVD (GannetLoad); GABA+ and Glx were fitted from the edited difference spectra (GannetFit); voxel tissue fractions (GM, WM, CSF) were obtained via SPM12-based co-registration and segmentation (GannetCoRegister/GannetSegment); and metabolite concentrations were expressed as tissue-corrected67 and α-corrected institutional-unit values, with 45% macromolecule contamination68 assumed for GABA+ (GannetQuantify). Spectra were retained only if SNR > 5, FWHM < 30 Hz, and fit error < 15%, evaluated independently for GABA+ and Glx per session, yielding a variable MRS subsample size (Supplementary Table S9). Resting-state BOLD data were preprocessed following a standard pipeline: slice-timing correction, T1 segmentation, distortion and motion correction using paired spin-echo field maps, coregistration, nuisance regression, MNI normalization, 6 mm smoothing, median filtering, and detrending.

#### Hurst estimation (WAND)

To accommodate both stationary and non-stationary dynamics within a unified framework, we employed an extended Hurst exponent parameterization supporting H up to 2. H was estimated using the maximum likelihood method implemented in the nonfractal MATLAB toolbox^25,69^, which models the averaged BOLD signal *X*_*t*_ at each parcel as a fractionally integrated noise (FIN) process^70^:

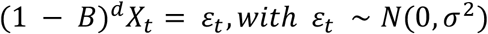

where *B* is the backshift operator and *d* is the fractional integration parameter governing long-memory properties. The Hurst exponent is related to *d* by H = d + 0.5. Parameters were estimated by minimizing the negative log-likelihood over the wavelet domain:

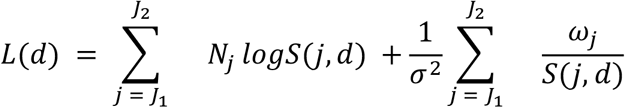

where *j* indexes wavelet scale, [J1,J2] is the range of scales included in the estimation, N_j_ is the number of wavelet coefficients at scale j, w_j_ is the observed wavelet variance at scale j, and S(j,d) is the theoretical spectral density of the FIN process at scale j.

Wavelet decomposition used the maximal overlap discrete wavelet transform (MODWT^71^) from the wmtsa MATLAB toolbox^72^ with a D4 filter, retaining levels j = [1, 5] (frequency bands: 0.008-0.25 Hz). MODWT was preferred over the standard DWT because it does not require truncation of the signal to the nearest power of 2, an important advantage given the limited time-series lengths in this study^73^. Bound parameters were lb = [-0.5, 0] and ub = [1.5, 10], yielding H ∈ [0, 2]. More details on Hurst estimation are reported in Supplementary Methods.

### Statistical analysis (WAND)

Because WAND’s MRS protocol was restricted to the resting state session, this discovery analysis is limited to associations between H and Glx/GABA+ in the absence of any cognitive stimulation; task-based modulation is addressed separately in the replication sample below. For each MRS voxel, we tested whether resting-state H within the spatially co-localized cortical parcels was modulated by the MRS-derived E/I index and its interaction with the structural hierarchy, fitting data jointly across all four voxels using linear mixed-effects models. After matching participants with both MRS spectra and resting-state H estimates, the final sample sizes were 150 for lSMN, 148 for ACC, 147 for OCC and 131 for rAUD voxels.

Before testing the cytoarchitectural models below, we first asked whether H and Glx/GABA+ simply differed in level across the four voxels. We conducted omnibus Friedman tests (Kendall’s W as effect size) on mean parcel-wise H and, separately, on Glx/GABA+ across the four voxels, followed by post-hoc pairwise Wilcoxon signed-rank tests (effect size r) with FDR correction across the six voxel pairs.

### Replication sample

#### Participants

MR data were collected from 36 healthy participants (19 males, 5 left-handed, age: 25.6 ± 3.2 years). Ethical approval for the study was obtained from the Ethical Committee of the University of Trento.

#### Experimental design

All data were acquired as part of a previously reported study^10^; full details of the experimental protocol and acquisition parameters were provided there. An interleaved fMRI-edited fMRS design was employed, beginning with a resting-state fMRI (rs-fMRI) scan serving as a baseline. Participants fixated on a central cross and were instructed to refrain from deliberate thought. Three subsequent task-based fMRI (tb-fMRI) runs administered an N-back working memory paradigm at increasing cognitive loads. In the 0-Back run, participants responded whenever a target stimulus appeared; each stimulus was displayed for 2 s, followed by a 0.5 s fixation cross. Each run comprised four blocks of 44 trials, including 9 congruent (target-matching) trials. The 1-Back run required participants to judge whether the current stimulus matched the immediately preceding one; the 2-Back run required comparison with the stimulus two trials prior. Block and trial structure were identical across runs. Stimuli were randomly drawn from numbers and letters throughout.

#### MR acquisition and preprocessing

MR acquisition and initial preprocessing followed the protocol of our prior publication. Data were acquired on a 3 T MAGNETOM Prisma (Siemens Healthcare) with a 64-channel head-neck coil. The protocol included: multiecho MPRAGE T1-weighted imaging (1 mm isotropic; TR = 2.5 s, TE = [1.69, 3.55, 5.41, 7.27] ms); four BOLD EPI sessions (3 mm isotropic; TR = 2 s, TE = 28 ms); double-echo field maps; and MEGA-PRESS fMRS (3 cm voxel; TE/TR = 68/2000 ms; 192 rest / 224 task averages) for GABA+ and Glx quantification. TR/TE, editing scheme, and voxel volumes closely matched the discovery (WAND) acquisition, enabling direct methodological replication. fMRS spectra were processed in Gannet (v3.2.0), following the same pipeline and quality-screening criteria described for the discovery sample above (see Discovery sample: MR Acquisition and Processing (WAND), above). fMRI preprocessing followed the same pipeline as the discovery (WAND) sample, with the exception of susceptibility distortion correction, performed using double-echo field maps, and of nuisance regression, which was applied only to the resting-state run.

For task-based runs, a general linear model (GLM) was estimated at the subject level in FSL to isolate task-evoked BOLD responses prior to estimating the Hurst exponent (H). Each run comprised four blocks (onsets: 0, 120, 240, 360 s; duration: 110.7 s) separated by ~9.3 s inter-block rest intervals. Three explanatory variables were modeled: sustained block activation, individual trial onsets (impulse; duration = 0 s), and congruent target trials, each convolved with a double-gamma hemodynamic response function^74^. GLM residuals representing BOLD variance unexplained by the task structure were extracted separately for each run (0-Back, 1-Back, 2-Back) and used as input to Hurst exponent estimation, ensuring that H reflects intrinsic neural dynamics rather than stimulus-locked activity^75^.

#### Eye-tracker acquisition and preprocessing

Eye-tracking data were recorded using an Eyelink 1000^76^ during all task-based runs. Pupil diameter signals were preprocessed as follows: (i) blink intervals were identified from discrete events in the EDF file, each expanded by a 100 ms margin ([start −0.1 s, end +0.1 s]), and affected samples were replaced using PCHIP interpolation^77^ between adjacent valid samples; (ii) outliers were detected from the absolute velocity signal using a threshold of median(speed) + 2·MAD^78^, and replaced via PCHIP interpolation; (iii) the cleaned signal was low-pass filtered with a zero-phase 4th-order Butterworth filter (cutoff: 9 Hz); (iv) the time series was decimated to 0.5 Hz using the signal.decimate function from SciPy^79^ (ftype=‘fir’, zero_phase=True); (v) the first sample (corresponding to the initial 2 s of acquisition) was discarded; (vi) subjects with invalid data or prominent artifacts upon visual inspection were excluded.

Microsaccade time series were extracted as follows: (i) onset and offset times were retrieved from EDF files; (ii) microsaccades with duration > 1 s were discarded; (iii) onset times were binned into 1 s bins; (iv) the resulting time series was decimated to 0.5 Hz as described above; (v) subjects excluded during pupil preprocessing were also excluded here.

Before Hurst exponent estimation, task-evoked components were regressed out of both signals using the same block-structure GLM employed for fMRI residualization (block activation, trial onsets, congruent target trials), fitted separately for each run. The residuals were retained for subsequent long-range dynamics analysis.

#### Hurst estimation

H was estimated using the same FIN-based maximum-likelihood framework and MODWT decomposition described for the discovery sample above (full derivation in Discovery sample: Hurst estimation (WAND), above).

The Hurst exponent was estimated on three distinct signal types. For the cortical analysis, H was computed parcel-wise on fMRI BOLD time series, separately for the resting-state run and for the GLM-residualized task runs (0-Back, 1-Back, 2-Back), yielding a spatial map of temporal scaling properties across the 100 parcels of the Schaefer atlas for each session. For the behavioral analyses, H was estimated on two eye-tracking signals, downsampled to the fMRI sampling rate to ensure temporal comparability and enable the same H-estimation pipeline: (i) GLM-residualized pupil diameter time series, reflecting slow arousal fluctuations; and (ii) GLM-residualized microsaccade rate time series, reflecting oculomotor suppression dynamics. In both cases, preprocessing and GLM residualization followed the procedures described above (Eye-Tracker Acquisition and Preprocessing), and H estimation used the same FIN-based maximum likelihood framework applied to the fMRI data.

### Statistical analysis

To characterize the spatial distribution of H across the cortex and identify regions with consistent group-level signal, parcel-wise mean H and standard deviation were computed across subjects separately for each session (rest, 0-Back, 1-Back, 2-Back) and rendered as cortical surface maps. To then identify cortical regions exhibiting reliable group-level deviations, voxel-wise one-sample t-tests were conducted for each condition using FSL randomise^80^ with 5000 permutations and Threshold-Free Cluster Enhancement (TFCE)^81^ correction, thresholded at p < 0.02 (1 - p_TFCE_ > 0.98), using a more stringent significance threshold to account for the four session-wise statistical tests.

To reduce dimensionality and isolate the dominant spatial patterns of H variation across subjects and sessions, singular value decomposition (SVD) was applied to a parcel × (subject·session) H matrix of size 100 × 140, centering by column mean. Missing values were imputed using the parcel-wise median across subjects, or the global median when unavailable. SVD factorizes X as:

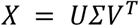

where U contains left singular vectors describing spatial distributions across parcels; Σ is a diagonal matrix of singular values reflecting variance explained; and *V*^*T*^ contains right singular vectors capturing how each spatial pattern is expressed across subjects and sessions. To assess the significance of the explained variance, a permutation test (2000 permutations) was performed by independently shuffling values within each column of X and recomputing the SVD. Empirical p-values were calculated as p = (q+1)/(P+1), where q is the number of permutations in which the permuted variance explained equaled or exceeded the observed value.

To formally test whether H is modulated by working memory load at each cortical location, parcel-wise repeated-measures ANOVAs were estimated with Session (4 levels: rest, 0-Back, 1-Back, 2-Back) as a within-subject factor, separately for each of the 100 parcels of the Schaefer atlas^82^. Significant session effects (p < 0.05, FDR^83^-corrected across parcels) were followed by post-hoc pairwise t-tests with multiple comparison correction. Effect sizes were quantified as partial η^2^ and summarized at the network level using the seven canonical functional networks of the Schaefer 100-parcel atlas (Visual, SomatoMotor, Dorsal Attention, Salience/Ventral Attention, Limbic, Control, Default Mode). To move from parcel-level findings to a functionally interpretable systems-level description, at the network level, parcel-wise H values were averaged within each network, and one-way Friedman tests with Session as the within-subject factor were estimated, followed by post-hoc pairwise Wilcoxon tests. Pairwise between-network H comparisons were also conducted non-parametrically within each session. To quantify session-dependent changes in inter-network H distance, pairwise ΔH was computed for each network pair, yielding a 7 × 7 antisymmetric matrix per session. A Friedman test assessed whether inter-network H differences varied significantly across sessions (p < 0.05, FDR-corrected across pairs), followed by post-hoc pairwise comparisons.

To test whether individual differences in MRS-derived E/I balance account for session-specific variation in H, linear mixed-effects (LME) models were estimated:

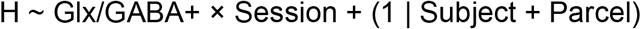

where Glx/GABA+ is the MRS-derived E/I index (Glx/GABA+ ratio), Session is a fixed categorical factor (4 levels), and Subject and Parcel are random intercepts. Models were estimated within the MRS voxel; two additional models decomposed Glx/GABA+ into GABA+ and Glx separately.

To evaluate the effects of N-back load and acquisition order on pupillometry-derived Hurst exponents, we fitted the following LME model:

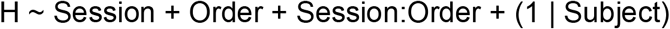

separately for H estimates derived from pupil size and microsaccade time series. Session denotes the n-back condition (0-Back, 1-Back, or 2-Back) while Order denotes the acquisition order (0 = fMRI 0-Back, 1 = fMRS 0-Back, 2 = fMRI 1-Back, 3 = fMRS 1-Back, 4 = fMRI 2-Back, and 5 = fMRS 2-Back).

To assess whether the association between neurometabolic balance and H also extended to pupillometry, we applied a model analogous to the one used for brain-derived H estimates to pupil-size and microsaccade time series. Because pupillometry measures provided one H value per subject and session, the parcel-level random effect was omitted:

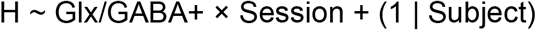

As in the brain analyses, additional models replaced the Glx/GABA+ ratio with GABA+ or Glx separately. All LME models were implemented in R using lme4 and lmerTest libraries.

Finally, to characterize the temporal dynamics of pupil diameter across task conditions and their relationship to Glx/GABA+ balance, and thereby probe an arousal-related physiological readout of the same neurometabolic axis, we used Growth Curve Analysis (GCA; Mirman, 2014), implemented as polynomial linear mixed-effects models predicting z-scored pupil diameter from orthogonal polynomial time terms (linear, quadratic, cubic), their interactions with working memory load and Glx/GABA+ ratio, and run order and scanner modality (fMRI/fMRS) as covariates. Three models were estimated. The first (GCA-1) tested whether cognitive load modulates trajectory shape:

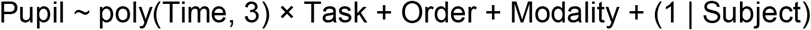

The second (GCA-2) tested whether Glx/GABA+ balance distorts the trajectory beyond task effects (Supplementary Materials):

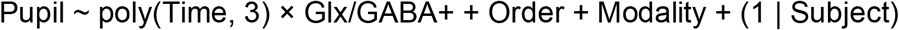

The third (GCA-3) tested whether the Glx/GABA+ effect varies with load, modeling a full three-way interaction:

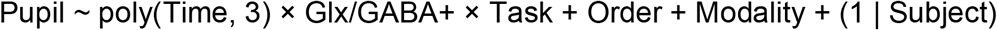

where Time is modeled as orthogonal polynomial terms (linear, quadratic, cubic), Task is working memory load, Glx/GABA+ is the MRS-derived E/I index, and run Order and scanner Modality (fMRI/fMRS) are included as covariates, with Subject as a random intercept. Group-mean observed trajectories were plotted alongside marginal fixed-effects predictions as a theoretical reference for the population-average waveform. All models were implemented in R using lme4 and lmerTest (see Supplementary Methods for full details).

### Statistical analysis (across cohorts)

We next tested whether the relationship between resting-state H and the MRS-derived E/I index varied along cortical cytoarchitecture on the discovery dataset. Parcels were defined as the Schaefer parcels overlapping each MRS volume, and each parcel’s cytoarchitectural position was characterized using two frameworks: discrete categories (von Economo atlas^44^) and a continuous microstructural axis (BigBrain gradient^45^). Each parcel’s hierarchical position was characterized by continuous predictors that were mean-centered to optimize interpretability and reduce polynomial multicollinearity.

Based on likelihood-ratio tests for optimal model selection, we implemented a second-order quadratic expansion for von Economo:

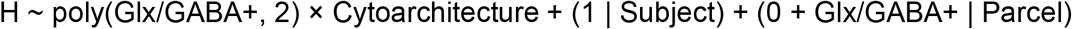

and a third-order cubic polynomial expansion for the continuous BigBrain gradient:

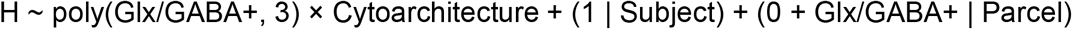

fit jointly across all four MRS voxels. To aid interpretation of the cubic interaction, we estimated the local slope of H with respect to Glx/GABA+ at the mean Glx/GABA+ level (mean-centered Glx/GABA+ = 0) at the agranular and granular ends of the observed BigBrain gradient using estimated marginal trends (emmeans/emtrends). Slopes were estimated at the minimum and maximum observed gradient values, and their difference was tested using pairwise contrasts with FDR correction. Two additional models replaced the Glx/GABA+ ratio with GABA+ or Glx separately, mirroring the discovery analysis.

Finally, to test whether H’s spatial organization reflects the intrinsic functional hierarchies that organize large-scale cortical activity, we related resting-state H to the principal gradient of macroscale cortical organization (Margulies et al., 2016) in both cohorts. Each Schaefer-100 parcel was assigned its group-average value from the published Schaefer-100 parcellation of this gradient, held constant across cohorts. We fitted linear mixed-effects models of the form:

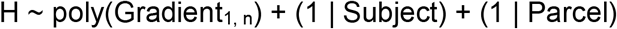

with n selected by nested model comparison (likelihood ratio tests) among linear, quadratic, and cubic specifications; models were fit by maximum likelihood in R using lme4 and lmerTest. This model was fitted separately at rest in the WAND cohort and, as a convergence check, in the replication sample. Because only the replication sample includes task data, we then asked whether this gradient-H relationship itself reorganizes under cognitive engagement, extending the model with an interaction between the gradient and session:

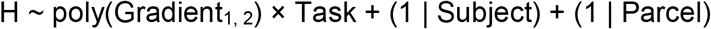

with Task as a four-level fixed factor (rest, 0-Back, 1-Back, 2-Back; rest as reference level) and the quadratic order carried over from the resting-state model above. To characterize how the linear and quadratic gradient terms varied across sessions, we followed the omnibus interaction test with session-specific simple-slope contrasts (estimated marginal trends, emmeans/emtrends), testing the significance of each polynomial term separately within rest, 0-Back, 1-Back and 2-Back, with correction for multiple comparisons across the four sessions. To characterise the spatial locus of this load-dependent change, we estimated the local slope of H on the principal gradient at representative unimodal (Gradient = −4) and transmodal (Gradient = +4) values, separately for each task condition. The difference between unimodal and transmodal slopes was tested formally within each task using pairwise contrasts, to assess whether the asymmetry in the Gradient 1-H relationship present at rest was preserved or attenuated under increasing cognitive load.

### Human Connectome Project dataset

To assess the reproducibility of the H topography across acquisition and preprocessing, we additionally analysed resting-state fMRI from the Human Connectome Project Young Adult dataset^39^: 100 unrelated participants with four runs each (two sessions, two phase-encoding directions runs; 1200 volumes per run, TR=0.72 s; 3T, multiband 8), taken from the minimally pre-processed release and parcellated with the Schaefer-100 atlas (cortical parcels only). Data underwent HCP minimal preprocessing, followed by regression of the linear trend, six motion parameters, white-matter and CSF signals and their derivatives; no band-pass filtering or global signal regression was applied. Time series were resampled to the TR of the discovery dataset (2 s) by polyphase rational resampling with an anti-aliasing FIR low-pass filter (MATLAB `resample`, factor 18/50; 1200 to 432 samples), so that wavelet scales j=1-5 covered the same frequency band (0.0078-0.25 Hz) as in the discovery cohort. Series were linearly detrended before resampling, and detrended and mean-centred again before estimation. H was estimated separately in each run with the same estimator and settings as in the other datasets and averaged across the four runs of each participant. Because absolute H depends on acquisition, preprocessing and series length, H values were z-scored across parcels within each dataset for display and network-level comparison.

## Author contributions

Matilde Bisi: Formal analysis, Methodology, Investigation, Software, Visualization, Writing - original draft; Asia Ferrari: Formal analysis, Data curation, Methodology, Investigation, Visualization, Software, Writing - original draft; J. Karolis Degutis: Methodology, Investigation, Software, Visualization, Writing - review & editing; Angela I. Renton: Methodology, Investigation, Writing - review & editing; Stefano Tambalo: Data curation, Investigation, Writing - review & editing; Michael V. Lombardo: Conceptualization, Investigation, Writing - review & editing; Jorge Jovicich: Conceptualization, Funding acquisition, Project administration, Investigation, Writing - review & editing; Dimitri Van De Ville: Funding acquisition, Investigation, Supervision, Writing - review & editing; Francesca Saviola: Conceptualization, Data curation, Formal analysis, Funding acquisition, Project administration, Investigation, Supervision, Visualization, Writing - original draft

## Acknowledgements

This work was supported in part by the Swiss National Science Foundation, grant No. 205321_207493 to F.S. and D.V.D.V., ISMRM Exchange Award 2021-2022 “Investigating in-vivo human brain dynamic connectivity with fast fMRI” to F.S. and Dipartimento di Eccellenza project 2018-2022 (Italian Ministry of Education, University and Research) to F.S.The replication dataset was acquired at University of Trento, Center for Mind/Brain Sciences]. We thank Dr. Ludovico Coletta, Alexandre Cionca, Chun Hei Michael Chan, Ekansh Sareen and Irene Bellin for their precious input and support during various phases of this study. Computing resources were provided by EPFL. The authors used Claude (Opus 5 and Opus 5.5; Anthropic) to check the manuscript for spelling and grammatical errors, formatting consistency, and clarity of language. No text, data, figures, or interpretations were generated by these tools. All output was reviewed by the authors, who take full responsibility for the content of this article.

## Data availability

The WAND dataset is publicly available via the GIN repository at [https://doi.org/10.1038/s41597-024-04154-7]. Data from the replication sample are available in processed and anonymized format in Zenodo (https://doi.org/10.5281/zenodo.15855362) or upon reasonable request to Prof. Jorge Jovicich. Data from the robustness sample (100-unrelated minimally preprocessed) are openly available as the WU-Minn Human Connectome Project (HCP) Young Adult dataset, through ConnectomeDB (https://db.humanconnectome.org).

## Code availability

The Hurst exponent estimation pipeline is available at https://github.com/wonsang/nonfractal. Analysis scripts reproducing adaptation of the Hurst estimation, all main figures and statistical results are available at https://github.com/MIPLabCH/EI. All custom code was written in MATLAB 9.13 and Python 3.11.

