## Supplementary Materials for "Cortical long-range dynamics track excitation-inhibition balance across cognitive states and organizational principles"

### Methods

#### MRS data quality control

Spectra were quality-screened separately for GABA+ and Glx in each session using the criteria described in the main Methods (SNR > 5, FWHM < 30 Hz, fit error < 15%), yielding a variable per-session MRS subsample. The number of participants retained per session and per metabolite is reported in Table S9; because quality control was applied independently to GABA+ and Glx, the analytic sample differs slightly between the composite MRS-derived E/I index and the single-metabolite models below. Quality screening was performed blind to task condition.

#### Hurst exponent estimation algorithm

The Hurst exponent estimation was performed using the maximum likelihood estimator implemented in bfn_mfin_ml from the nonfractal MATLAB toolbox[1,2]. As described in the main Methods, each time series was modeled as a fractionally integrated noise process[3], and the Hurst exponent was obtained from the fractional integration parameter as H=d+0.5.

For a given value of d, the theoretical scale-dependent variance of the FIN process at wavelet scale j was computed as:

$$S(j,d) \ = \ 2B_1(d)(2\pi)^{-2d}2^{-2dj}[1 + (2\pi)^2 \frac{d}{12}\frac{B_3(d)}{B_1(d)}]$$

where

$$B_1(d) \ = \ \frac{1 - 2^{2d-1}}{1-2d} \qquad\qquad B_3(d) \ = \ \frac{1 - 2^{2d-3}}{3-2d}$$

The fractional integration parameter was estimated by minimizing the wavelet-domain negative log-likelihood[4,1]:

$$L(d) \ = \ \sum_{j\,=\,J_1}^{J_2} \quad N_j \, logS(j,d) \ + \frac{1}{\sigma^2}\sum_{j\,=\,J_1}^{J_2} \quad \frac{\omega_j}{S(j,d)}$$

where $j$ indexes wavelet scale[4], $[J_1, J_2]$ = [1, 5] is the range of scales included in the estimation, $N_j$ is the number of wavelet coefficients at scale j, $w_j$ is the observed

wavelet variance at scale j, $\sigma^2$ is the variance of the fractionally integrated noise and S(j,d) is the theoretical spectral density of the FIN process at scale j.
The optimization was performed with an iterative algorithm of two steps:

1. First, the current value of $\sigma^2$ is calculated as a function of $d$:

$$\sigma^2 = \frac{var(X)}{2B_1(d)(2\pi)^{1-2d}\sum_{j=J_1}^{J2} \quad 2^{(2d-1)j}}$$

2. Then, the log-likelihood for the current $\sigma^2$ is minimized with respect to $d$.

The algorithm terminates when the relative change of $\sigma^2$ is lower than abstol, set to $10^{-10}$. The initial value of d was kept at the default value (0.1).

##### Glx/GABA+ and H within-voxel correlations

Pearson and Spearman correlations between Glx/GABA+ and cortical H were computed separately for each of the four WAND voxels (ACC, OCC, rAUD, and lSMN) and for the lDLPFC voxel in the replication dataset. For each subject, voxel-level H was calculated by averaging H across the cortical parcels overlapping with the corresponding MRS voxel.

##### Neurometabolic decomposition of cortical H

To test whether the composite Glx/GABA+ index or its constituent neurometabolites better explain session-dependent modulation of cortical H, GABA+ and Glx were each entered separately as predictors in place of the composite Glx/GABA+ ratio: H ~ GABA × Session + (1 | Subject + Parcel) and H ~ Glx × Session + (1 | Subject + Parcel).

##### Pupillometry- and microsaccade-derived H: session and order effects

Session and acquisition-order effects on pupil- and microsaccade-derived H were tested using H ~ Session + Order + Session:Order + (1 | Subject), fitted separately for each signal (see main Methods for the definition of Order).

##### Pupillometry- and microsaccade-derived H: session and modality effects

Because acquisition modality followed a fixed order across all participants and sessions (fMRI first, followed by fMRS), the second acquisition also represented a repeated exposure to the same cognitive-load condition. To assess whether pupil- and microsaccade-derived H differed between the first and second task repetition, and whether this effect varied across sessions, we fitted the model H ~ Session × Modality + (1 | Subject) separately for each signal.

##### Growth Curve Analysis: model specification

Epochs spanning −4 to 110 s relative to block onset were extracted for each of the four blocks per run, across six runs (0-Back fMRI, 0-Back fMRS, 1-Back fMRI, 1-Back fMRS, 2-Back fMRI, 2-Back fMRS). Time within each epoch was normalized across the full dataset, and three orthogonal polynomial regressors were derived. The linear term (ot1) captures the overall directional trend of the pupil response, reflecting sustained increases or decreases in arousal over time. The quadratic term (ot2) captures trajectory curvature, the characteristic initial constriction at task onset followed by partial recovery as attentional resources are recruited and stabilized. The cubic term (ot3) captures asymmetric late inflections such as secondary deflections reflecting fatigue-related fluctuations in arousal regulation during sustained effort. This three-degree polynomial was chosen based on the established pupil response profile during sustained cognitive effort, comprising an initial decline, a sustained trough, and a partial asymmetric recovery5. Interactions between these time terms and Task or MRS-derived E/I therefore test whether working memory load or individual Glx/GABA+ balance alters the overall directional trend (ot1 interaction), the sharpness and timing of the trough (ot2 interaction), or the late recovery dynamics (ot3 interaction) of the pupil trajectory.

Three models were fitted. GCA-1 tested the effect of cognitive load on the pupil trajectory (polynomial time × Task). GCA-2 tested the main effect of static Glx/GABA+ on trajectory shape, collapsed across tasks (polynomial time × Glx/GABA+). GCA-3 tested whether the relationship between Glx/GABA+and trajectory shape depends on load (polynomial time × Glx/GABA+ × Task). 0-Back was the reference condition. GCA-2 and GCA-3 were restricted to participants with eligible fMRS data.

Pupil diameter was z-scored within each participant before modeling; static (session-averaged) MRS-derived E/I indices (Glx/GABA+, GABA+, Glx) were z-scored across participants. Given the high temporal resolution of the pupil signal (~10 Hz; ~900,000 observations), models were fitted on data thinned by a factor of 10 (every 10th timepoint within each subject/task/block series); full-resolution data were used for visualization only. Random effects structures were selected by iterative simplification from the maximal converging model, retaining the most complex non-singular solution; random intercepts and slopes for the linear polynomial term over subjects were included where supported. Fixed effects were tested using Satterthwaite-approximated degrees of freedom6,7. Models were estimated using restricted maximum likelihood with a bobyqa optimizer.

##### MRS-derived E/I and peripheral H

The same H ~ Glx/GABA+ × Session linear mixed-effects framework used for cortical H was applied to the scalar per-session Hurst summaries of pupil and microsaccade signals, omitting the parcel-level random effect.

##### Task-dependent reconfiguration of cortical H: voxel-wise statistics

For each session (Rest, 0-Back, 1-Back, 2-Back), voxel-wise one-sample t-tests against zero were conducted using FSL randomise[8] (5,000 permutations, threshold-free cluster enhancement, $p < 0.02$). Post-hoc pairwise t-tests between all session pairs were then conducted at each parcel to characterise rest-task and task-load differences (see main Methods for full model specification).

##### Network-level statistics of H across sessions

Parcel-wise H values were averaged within each of the seven Yeo-Krienen networks[9]. For each network, a one-way Friedman test across the four sessions was followed by post-hoc pairwise Wilcoxon signed-rank tests (within-network tests). The same procedure was repeated within each session across all network pairs (within-session tests), and on the session-wise ΔH between every network pair (ΔH matrix tests). All omnibus and pairwise p-values were FDR-corrected.

##### Alternative parcellation (Glasser360) replication

To assess the robustness of our findings to parcellation choice, we replicated the main task-related Hurst exponent analyses using time series extracted with the Glasser360 atlas[10] in both the discovery and replication datasets.

##### Cytoarchitectural voxel composition

Von Economo class follows the five-class structural typology[11]: 1 = agranular, 2 = frontal, 3 = parietal, 4 = polar, 5 = granular, ordered from the thick, weakly laminated pole to the thin, granular-layer-IV-dominant primary sensory pole. BigBrain gradient values index each parcel's position along the continuous microstructural hierarchy described in the main text, with more negative values corresponding to the sensory-motor pole and more positive values to the associative pole.

#### Supplementary results

##### MRS data quality control

| | **GABA+** | **Glx** | **Both** |
|---|---|---|---|
| **Rest** | 23 | 20 | 20 |
| **0-Back** | 21 | 19 | 18 |
| **1-Back** | 22 | 21 | 20 |
| **2-Back** | 18 | 18 | 16 |

**Table S1.** Number of participants passing MRS quality control in each subsample, by neurometabolite (GABA+, Glx) and session.

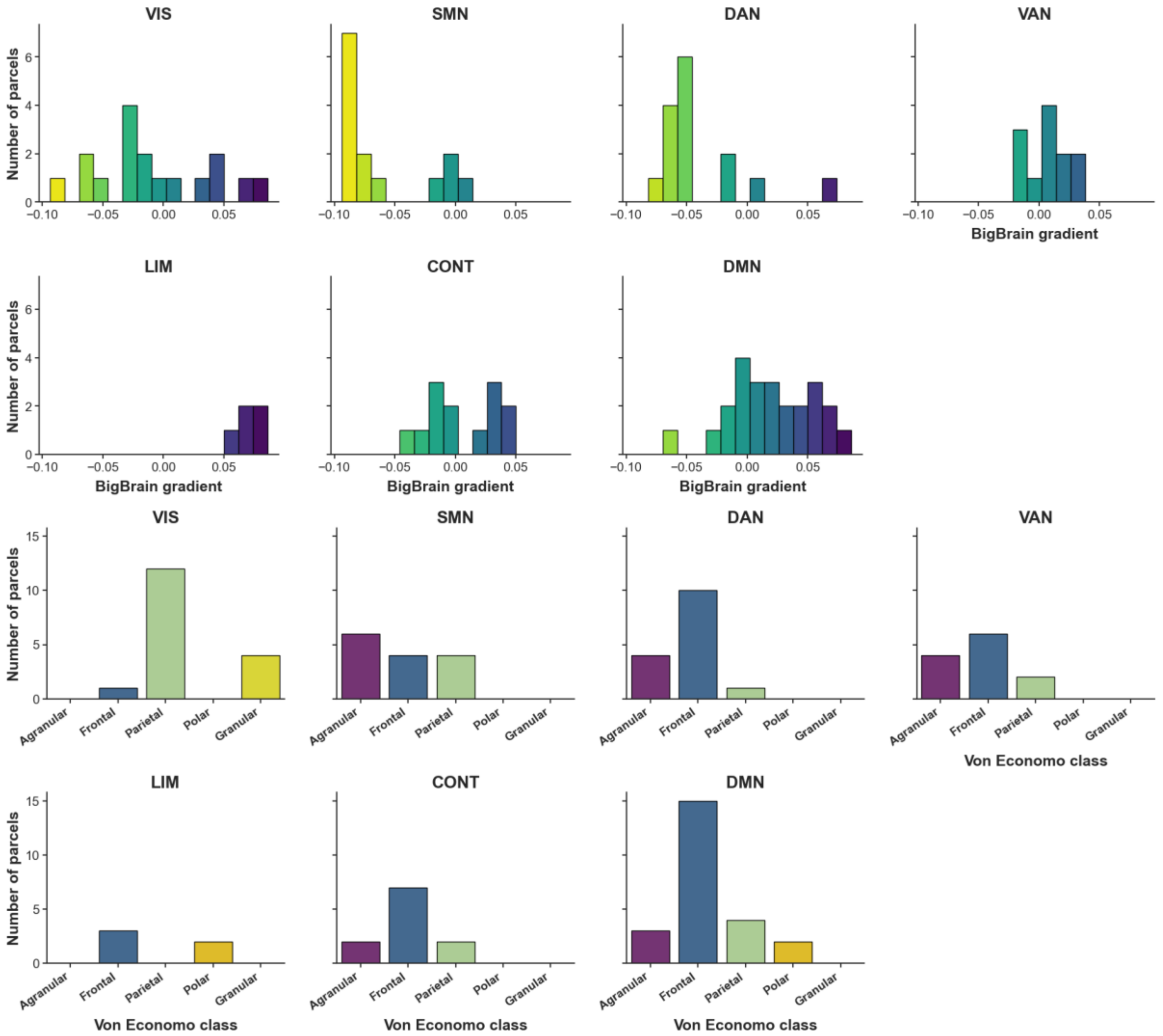


**Figure S1.** Distribution of BigBrain gradient values and Von Economo cytoarchitectonic classes across functional networks. Abbreviations: VIS, Visual; SMN, Somatomotor; DAN, Dorsal Attention; VAN, Salience/Ventral Attention; LIM, Limbic; CONT, Control; DMN, Default Mode Network.

#### MRS-derived E/I and H differences across voxels

Post hoc Wilcoxon tests between voxels showed significant differences in H for ACC−OCC ($r=0.84$, $\Delta$median=-0.20), ACC−lSMN ($r=0.75$, $\Delta$median=-0.18), OCC−rAUD ($r=0.80$, $\Delta$median=0.16), and rAUD−lSMN ($r=0.77$, $\Delta$median=-0.17); all $p_{FDR}<0.001$. MRS-derived E/I showed a largely opposite pattern, with significant differences for ACC−OCC ($r=0.86$, $\Delta$median=0.80), ACC−rAUD ($r=0.76$, $\Delta$median=0.35), ACC−lSMN ($r=0.87$, $\Delta$median=0.87), OCC−rAUD ($r=0.79$, $\Delta$median=-0.47), and rAUD−lSMN ($r=0.78$, $\Delta$median=0.51); all $p_{FDR}<0.001$.

#### Glx/GABA+ and H within-voxel correlation

None of the WAND voxels showed a significant Pearson or Spearman correlation between Glx/GABA+ and voxel-level H (Figure S1; ACC: $r = 0.048$, $p = 0.566$, $\rho = 0.072$, $p = 0.382$; OCC: $r = −0.015$, $p = 0.854$, $\rho = −0.026$, $p = 0.758$; rAUD: $r = −0.009$, $p = 0.916$, ρ = −0.015, p = 0.867; lSMN: r = −0.039, p = 0.640, ρ = −0.044, p = 0.597; lDLPFC: r = 0.011, p = 0.958, ρ = 0.081, p = 0.707).

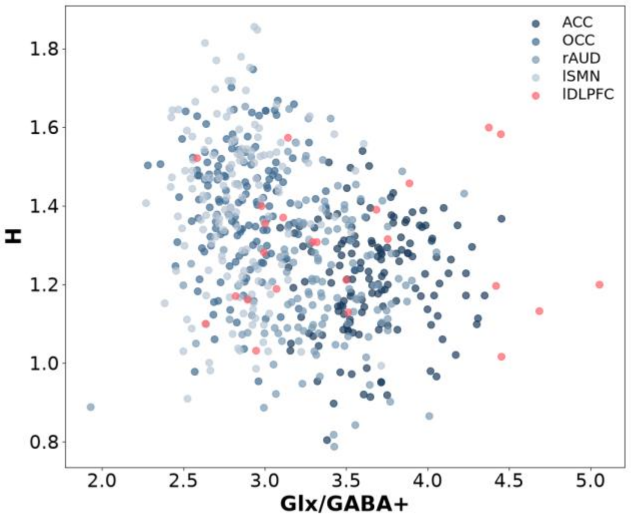


**Figure S2.** Scatterplot of the relationship between Glx/GABA+ and H within each MRS voxel. Each point represents a voxel of one participant.

##### Neurometabolic decomposition of cortical H

The linear mixed-effects models of the form H ~ GABA × Session + (1 | subject + parcel) and H ~ Glx × Session + (1 | subject + parcel) showed no significant association between GABA or Glx and H in the Rest condition (GABA: $\beta = 0.09$, $p_{FDR} = 0.06$, Glx: $\beta = -0.01$, $p_{FDR} = 0.702$). The Session interaction terms were also non-significant (GABA 0-Back: $\beta = -0.10$, $p_{FDR} = 0.149$; 1-Back: $\beta = -0.11$, $p_{FDR} = 0.088$; 2-Back: $\beta = -0.05$, $p_{FDR} = 0.530$, Glx 0-Back: $\beta = 0.01$, $p_{FDR} = 0.718$; 1-Back: $\beta = 0.01$, $p_{FDR} = 0.708$; 2-Back: $\beta = 0.04$, $p_{FDR} = 0.105$).

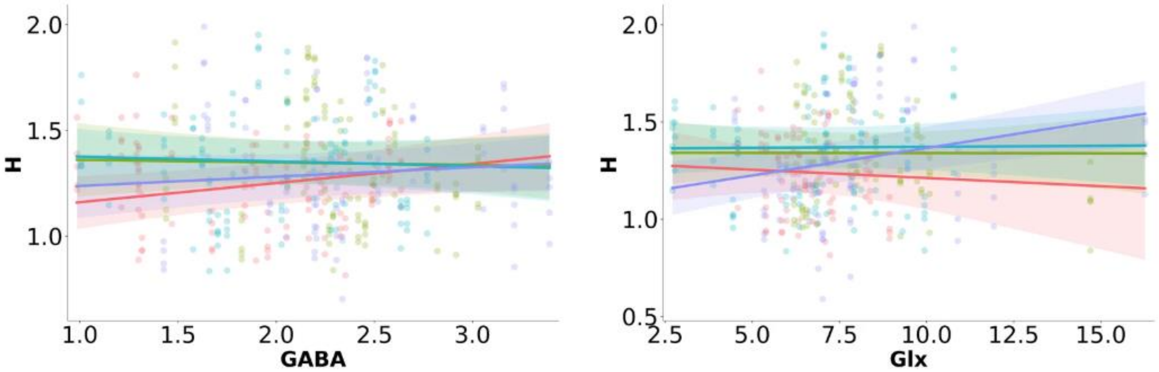


**Figure S3.** Relationship between cortical H and GABA+ (left) or Glx (right), fit separately for each session.

##### Pupillometry-derived H

Hurst exponents estimated on GLM-residual pupil diameter signals at the fMRI-matched timescale (0.5 Hz, 0.008–0.25 Hz band) revealed a significant effect of run order ($\beta = -0.09$, $p = 0.013$), reflecting a gradual decrease in H across successive runs consistent with arousal habituation over the session. This order effect was further qualified by a significant task × order interaction in the 1-Back condition ($\beta = +0.11$, $p = 0.025$). Because of this interaction, the task effect on H is best interpreted as order-dependent rather than constant; averaged across runs, neither the 1-Back nor the 2-Back condition produced a significant marginal effect relative to rest (1-Back: $\beta = -0.15$, $p = 0.24$; 2-Back: $\beta = +0.03$, $p = 0.87$), but this average masks the run-dependent modulation revealed by the interaction.

##### Microsaccade-derived H

Hurst exponents of microsaccade rate time series showed stronger and more consistent modulation by cognitive load than pupil diameter. A significant main effect of task was found for 1-Back ($\beta = -0.28$, $p < 0.001$), reflecting reduced temporal persistence of microsaccadic dynamics under moderate load relative to 0-Back; the 2-Back condition did not differ significantly from baseline ($\beta = +0.007$, $p = 0.95$). Run order exerted a strong negative effect on H ($\beta = -0.14$, $p < 0.001$), indicating progressive suppression of fractal dynamics across the session. Significant task × order interactions were found for both 1-Back ($\beta = +0.13$, $p < 0.001$) and 2-Back ($\beta = +0.07$, $p = 0.02$), suggesting that cognitive load partially counteracts the order-driven H decline.

##### Pupillometry- and microsaccade-derived H: session and modality effects

For pupil-derived H, fMRS acquisition was associated with significantly lower H than fMRI in the reference 0-Back condition ($\beta = -0.086$, $p = 0.013$). This difference was significantly attenuated in the 1-Back condition, as indicated by a positive Session × Modality interaction ($\beta = +0.107$, $p = 0.025$), whereas no significant interaction was observed for 2-Back ($\beta = +0.050$, $p = 0.294$). A similar but stronger pattern was observed for microsaccade-derived H. H was significantly lower during fMRS than fMRI acquisition in the reference 0-Back condition ($\beta = -0.135$, $p < 0.001$), with significant positive interactions for both 1-Back ($\beta = +0.128$, $p < 0.001$) and 2-Back ($\beta = +0.067$, $p = 0.025$). Thus, the reduction in H associated with the second acquisition was strongest during 0-Back and was attenuated under higher cognitive load.

##### Growth curve analysis:

During 0-Back, pupil responses showed a strong negative linear trend and a positive quadratic component (initial constriction followed by partial recovery), with an asymmetric late inflection (cubic term; all $p < 0.001$). Linear and quadratic components were significantly modulated by load ($p < 0.001$): 1-Back and 2-Back attenuated the negative linear decline, while 1-Back selectively deepened the quadratic trough ($\beta = +54.6$, $p < 0.001$) but 2-Back did not ($p > 0.05$), indicating load-dependent differences in trough depth and timing (Figure 2G). The main-effect model (GCA-2) estimated the influence of MRS-derived E/I balance on pupil trajectory shape collapsed across tasks, in the 22 participants with eligible fMRS spectroscopy data. The most complex non-singular random-effects structure retained random intercepts and a random slope for the linear polynomial term over subjects. EIB exerted a significant positive main effect on mean pupil diameter ($F_{(1,2997)} = 26.1$, $p < 0.001$; $\beta = +0.039$), indicating that higher cortical excitatory tone was associated with larger overall pupil responses across task conditions, and significantly modulated all three polynomial components of the pupil trajectory: the linear term ($F_{(1,785)} = 6.2$, $p = 0.013$; $\beta = +16.7$), reflecting attenuation of the overall negative temporal trend with increasing Glx/GABA+; the quadratic term ($F_{(1,47260)} = 13.2$, $p < 0.001$; $\beta = +15.8$), reflecting a more pronounced mid-epoch trough; and the cubic term ($F_{(1,47260)} = 4.7$, $p = 0.030$; $\beta = +9.5$), reflecting steeper late asymmetric recovery dynamics. These results indicate that individual Glx/GABA+ balance shapes both the amplitude and the temporal structure of the sustained arousal response, independent of working memory load. Allowing the Glx/GABA+ effect to vary by task, significant three-way Glx/GABA+ × Task × polynomial-component interactions emerged for all components (all $p < 0.05$; Figure 2H), driven mainly by 1-Back, where Glx/GABA+ predicted the linear decline, the mid-epoch trough and the late recovery (all $p < 0.05$). Under 2-Back, Glx/GABA+ no longer shaped trajectory structure but produced a sustained elevation in overall pupil diameter ($\beta = +0.11$, $p < 0.001$). Glx/GABA+'s expression in arousal dynamics therefore shifts from temporal structure to tonic level as cognitive demand increases, refining the load-independent main effect reported above (GCA-2).

##### MRS-derived E/I and peripheral H

We applied the same H ~ Glx/GABA+ × Session linear mixed-effects framework used for cortical H (see main text) to test whether MRS-derived E/I balance predicts the scalar Hurst summaries of pupil and microsaccade signals across sessions (Figure S3). No significant relationships emerged between either Hurst exponent and MRS-derived E/I balance, whether modelled as the composite Glx/GABA+ ratio or decomposed into GABA+ and Glx separately, nor were there significant interactions with session. For microsaccade H, no equivalent time-resolved model of Glx/GABA+ modulation was tested, so these data cannot determine whether peripheral microsaccadic persistence is genuinely insensitive to Glx/GABA+ or whether the null result instead reflects reduced statistical power.

This null result contrasts with the significant Glx/GABA+-by-load effects recovered by the growth curve analysis of pupil diameter (see “Growth curve analysis: main-effect Glx/GABA+ model (GCA-2)”, below), and more plausibly reflects a difference in analytical sensitivity than a genuine absence of coupling between prefrontal Glx/GABA+ and peripheral fractal dynamics: the GCA models retain the full time-resolved signal from the fMRS-eligible subsample, whereas the present models compress each subject's session-long peripheral time series into a single scalar Hurst estimate per session, discarding the temporal structure to which GCA is sensitive.

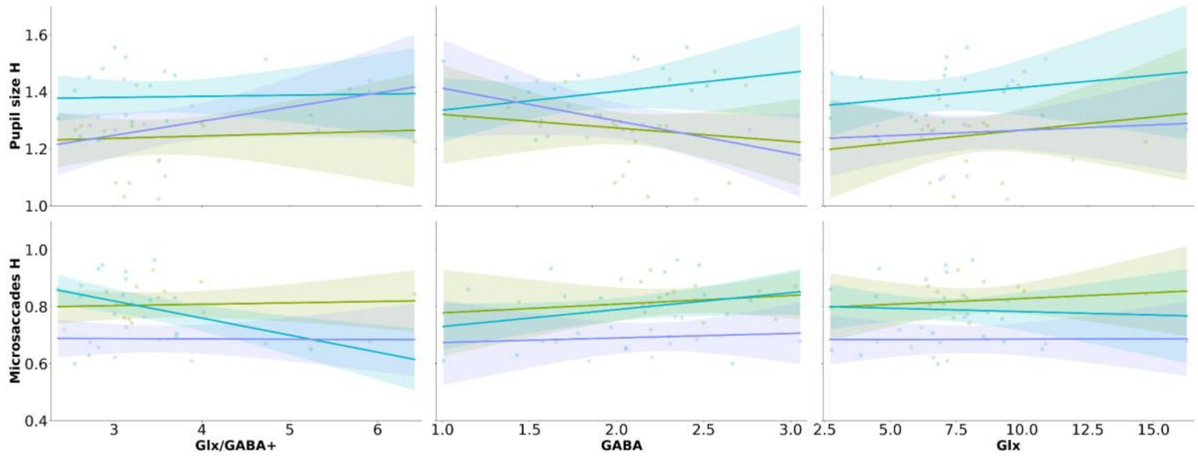


**Figure S4.** Relationship between pupil- and microsaccades derived H for Glx/GABA+, GABA+ and Glx, fit separately for each session.

#### Task-dependent reconfiguration of cortical H: voxel-wise statistics

One-sample t-tests performed separately for each session revealed that H was significantly greater than zero across large portions of the cortex in all four sessions (TFCE-corrected, $p<0.02$). The spatial distribution of t-statistics was broadly similar across sessions, with high values distributed across the cortical surface rather than concentrated in a single focal cluster. Percentile maps confirmed that the magnitude of the effect was spatially heterogeneous: the top 20% of parcels showed distributed clusters of stronger t-statistics, whereas the bottom 20% showed relatively weaker effects. H exceeded zero in the majority of cortical parcels in every session, with heterogeneous effect magnitude across regions.

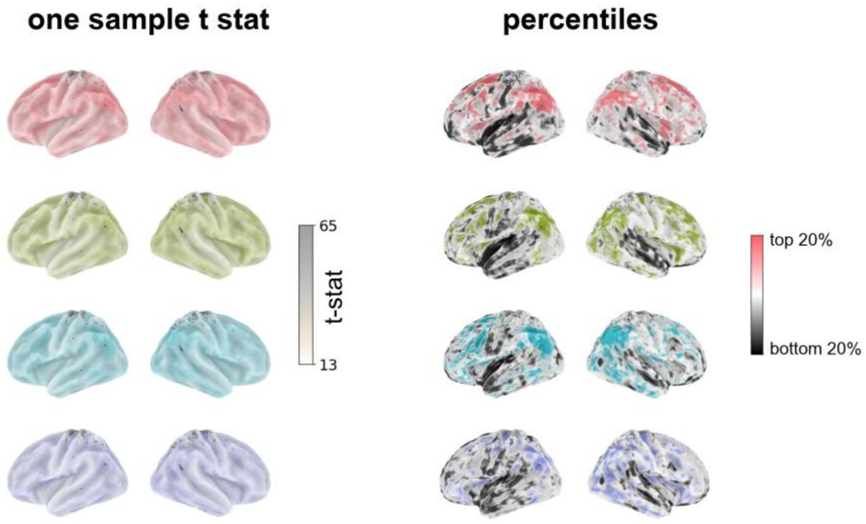


**Figure S5.** Voxel-wise one-sample t-statistics for H > 0 in each session (left) and percentile maps of effect magnitude (right).

#### Pairwise comparisons between all session pairs

Post-hoc pairwise t-tests between sessions revealed spatially heterogeneous changes in H (Figure S5 and Table S2). The rest–task differences followed a region-dependent pattern, with H tending to increase in frontal and temporal regions and decrease in a smaller number of parietal parcels. Spatial patterns were broadly similar for the rest vs 0-Back and rest vs 1-Back contrasts, with prominent involvement of the limbic, control, and default-mode networks; the rest vs 2-Back contrast additionally involved parcels in the somatomotor, dorsal-attention, and salience ventral-attention networks. Task-load modulation showed a decrease in H with increasing working-memory load across several parcels, most evident in the default-mode network, followed by somatomotor, limbic, salience ventral-attention, and control regions. This effect was absent between 0-Back and 1-Back, but became more pronounced in the 0-Back vs 2-Back and 1-Back vs 2-Back contrasts.

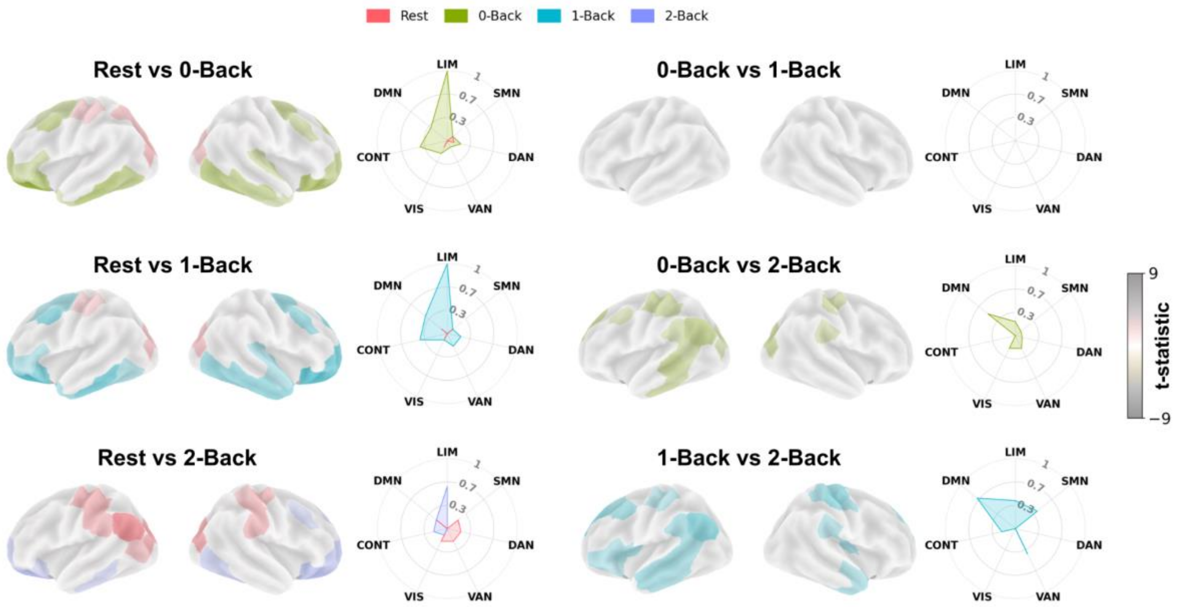


**Figure S6.** Post-hoc pairwise between-session t-statistic maps (left, per comparison) and network-averaged radar plots of significant differences (right).

| Parcel | Network | Pairwise comparison | t | p_FDR | N |
|---|---|---|---|---|---|
| 1 | Vis | Rest vs 0-Back | -3.26 | 0.0158 | 33 |
| 1 | Vis | 0-Back vs 2-Back | 2.54 | 0.0482 | 33 |
| 7 | Vis | Rest vs 2-Back | 3.09 | 0.0241 | 34 |
| 8 | Vis | Rest vs 0-Back | 3.59 | 0.0032 | 34 |
| 8 | Vis | Rest vs 1-Back | 3.08 | 0.0084 | 34 |
| 8 | Vis | Rest vs 2-Back | 4.48 | 5.08e-04 | 34 |
| 8 | Vis | 0-Back vs 2-Back | 2.49 | 0.0272 | 34 |
| 14 | SomMot | Rest vs 0-Back | 2.46 | 0.0305 | 30 |
| 14 | SomMot | Rest vs 1-Back | 2.30 | 0.0349 | 30 |
| 14 | SomMot | Rest vs 2-Back | 3.92 | 0.0030 | 30 |
| 14 | SomMot | 0-Back vs 2-Back | 3.18 | 0.0104 | 30 |
| 14 | SomMot | 1-Back vs 2-Back | 2.50 | 0.0305 | 30 |

| | | | | | |
|---|---|---|---|---|---|
| 16 | DorsAttn | Rest vs 0-Back | -3.85 | 0.0034 | 32 |
| 16 | DorsAttn | Rest vs 1-Back | -2.77 | 0.0280 | 32 |
| 17 | DorsAttn | Rest vs 2-Back | 3.63 | 0.0067 | 29 |
| 18 | DorsAttn | Rest vs 0-Back | 2.90 | 0.0426 | 29 |
| 19 | DorsAttn | Rest vs 2-Back | 3.90 | 0.0031 | 30 |
| 19 | DorsAttn | 0-Back vs 2-Back | 2.87 | 0.0230 | 30 |
| 23 | DorsAttn | Rest vs 0-Back | -3.02 | 0.0148 | 33 |
| 23 | DorsAttn | Rest vs 1-Back | -3.03 | 0.0148 | 33 |
| 24 | SalVentAttn | Rest vs 2-Back | 3.38 | 0.0116 | 33 |
| 24 | SalVentAttn | 0-Back vs 2-Back | 2.41 | 0.0434 | 33 |
| 24 | SalVentAttn | 1-Back vs 2-Back | 3.02 | 0.0150 | 33 |
| 26 | SalVentAttn | 1-Back vs 2-Back | 3.10 | 0.0236 | 34 |
| 30 | SalVentAttn | Rest vs 0-Back | -3.34 | 0.0062 | 34 |
| 30 | SalVentAttn | Rest vs 1-Back | -3.78 | 0.0038 | 34 |
| 30 | SalVentAttn | 1-Back vs 2-Back | 2.81 | 0.0167 | 34 |
| 31 | Limbic | Rest vs 0-Back | -7.99 | 2.42e-08 | 33 |
| 31 | Limbic | Rest vs 1-Back | -6.61 | 5.61e-07 | 33 |
| 31 | Limbic | Rest vs 2-Back | -4.68 | 1.01e-04 | 33 |
| 32 | Limbic | Rest vs 0-Back | -2.65 | 0.0367 | 34 |
| 32 | Limbic | Rest vs 1-Back | -3.25 | 0.0158 | 34 |
| 33 | Limbic | Rest vs 0-Back | -4.79 | 1.04e-04 | 34 |
| 33 | Limbic | Rest vs 1-Back | -5.07 | 9.13e-05 | 34 |
| 33 | Limbic | Rest vs 2-Back | -2.60 | 0.0166 | 34 |
| 33 | Limbic | 0-Back vs 2-Back | 3.07 | 0.0085 | 34 |
| 33 | Limbic | 1-Back vs 2-Back | 2.87 | 0.0108 | 34 |
| 37 | Cont | Rest vs 0-Back | -2.74 | 0.0205 | 31 |
| 37 | Cont | Rest vs 1-Back | -3.11 | 0.0147 | 31 |
| 37 | Cont | 1-Back vs 2-Back | 3.04 | 0.0147 | 31 |
| 38 | Default | 1-Back vs 2-Back | 3.20 | 0.0189 | 32 |
| 39 | Default | 0-Back vs 2-Back | 3.77 | 0.0040 | 33 |
| 39 | Default | 1-Back vs 2-Back | 3.35 | 0.0062 | 33 |
| 40 | Default | 0-Back vs 2-Back | 2.89 | 0.0201 | 34 |
| 40 | Default | 1-Back vs 2-Back | 3.15 | 0.0201 | 34 |
| 41 | Default | Rest vs 2-Back | 6.20 | 8.80e-06 | 27 |
| 41 | Default | 0-Back vs 2-Back | 4.00 | 9.23e-04 | 27 |
| 41 | Default | 1-Back vs 2-Back | 4.34 | 5.78e-04 | 27 |
| 42 | Default | Rest vs 0-Back | -4.54 | 2.13e-04 | 34 |
| 42 | Default | Rest vs 1-Back | -4.65 | 2.13e-04 | 34 |
| 42 | Default | Rest vs 2-Back | -2.70 | 0.0218 | 34 |
| 42 | Default | 1-Back vs 2-Back | 2.32 | 0.0404 | 34 |
| 43 | Default | Rest vs 0-Back | -3.02 | 0.0271 | 34 |
| 43 | Default | Rest vs 1-Back | -2.67 | 0.0271 | 34 |
| 43 | Default | 1-Back vs 2-Back | 2.61 | 0.0271 | 34 |
| 44 | Default | Rest vs 2-Back | 3.39 | 0.0039 | 32 |
| 44 | Default | 0-Back vs 2-Back | 3.80 | 0.0038 | 32 |
| 44 | Default | 1-Back vs 2-Back | 3.39 | 0.0039 | 32 |
| 45 | Default | Rest vs 0-Back | -3.88 | 0.0020 | 32 |
| 45 | Default | Rest vs 1-Back | -3.78 | 0.0020 | 32 |
| 45 | Default | Rest vs 2-Back | -2.52 | 0.0340 | 32 |
| 46 | Default | 0-Back vs 2-Back | 3.67 | 0.0029 | 30 |
| 46 | Default | 1-Back vs 2-Back | 4.08 | 0.0019 | 30 |

| | | | | | |
|---|---|---|---|---|---|
| 47 | Default | Rest vs 0-Back | -3.86 | 0.0016 | 33 |
| 47 | Default | Rest vs 1-Back | -4.39 | 6.91e-04 | 33 |
| 47 | Default | 0-Back vs 2-Back | 2.29 | 0.0435 | 33 |
| 47 | Default | 1-Back vs 2-Back | 3.36 | 0.0041 | 33 |
| 48 | Default | Rest vs 0-Back | -2.41 | 0.0453 | 30 |
| 48 | Default | Rest vs 1-Back | -2.56 | 0.0453 | 30 |
| 48 | Default | 1-Back vs 2-Back | 3.05 | 0.0294 | 30 |
| 49 | Default | Rest vs 1-Back | 2.28 | 0.0442 | 32 |
| 49 | Default | Rest vs 2-Back | 4.74 | 2.68e-04 | 32 |
| 49 | Default | 0-Back vs 2-Back | 4.24 | 5.57e-04 | 32 |
| 49 | Default | 1-Back vs 2-Back | 2.98 | 0.0112 | 32 |
| 50 | Default | Rest vs 0-Back | 2.22 | 0.0405 | 33 |
| 50 | Default | Rest vs 1-Back | 2.64 | 0.0192 | 33 |
| 50 | Default | Rest vs 2-Back | 5.12 | 8.52e-05 | 33 |
| 50 | Default | 0-Back vs 2-Back | 4.44 | 3.00e-04 | 33 |
| 50 | Default | 1-Back vs 2-Back | 3.84 | 0.0011 | 33 |
| 51 | Vis | Rest vs 0-Back | -4.53 | 3.27e-04 | 32 |
| 51 | Vis | Rest vs 1-Back | -4.43 | 3.27e-04 | 32 |
| 51 | Vis | Rest vs 2-Back | -3.65 | 0.0019 | 32 |
| 53 | Vis | Rest vs 0-Back | -4.49 | 4.93e-04 | 34 |
| 53 | Vis | Rest vs 1-Back | -3.67 | 0.0025 | 34 |
| 53 | Vis | Rest vs 2-Back | -2.88 | 0.0140 | 34 |
| 57 | Vis | Rest vs 0-Back | 2.64 | 0.0189 | 33 |
| 57 | Vis | Rest vs 1-Back | 2.96 | 0.0115 | 33 |
| 57 | Vis | Rest vs 2-Back | 4.61 | 3.67e-04 | 33 |
| 57 | Vis | 0-Back vs 2-Back | 3.25 | 0.0081 | 33 |
| 59 | SomMot | Rest vs 0-Back | -2.93 | 0.0208 | 34 |
| 59 | SomMot | Rest vs 1-Back | -2.88 | 0.0208 | 34 |
| 60 | SomMot | 1-Back vs 2-Back | 3.11 | 0.0233 | 33 |
| 63 | SomMot | 1-Back vs 2-Back | 3.14 | 0.0214 | 34 |
| 64 | SomMot | Rest vs 2-Back | 3.44 | 0.0048 | 34 |
| 64 | SomMot | 0-Back vs 2-Back | 3.02 | 0.0096 | 34 |
| 64 | SomMot | 1-Back vs 2-Back | 4.01 | 0.0019 | 34 |
| 65 | SomMot | Rest vs 2-Back | 2.64 | 0.0392 | 30 |
| 65 | SomMot | 1-Back vs 2-Back | 3.05 | 0.0294 | 30 |
| 66 | SomMot | 1-Back vs 2-Back | 3.30 | 0.0141 | 34 |
| 68 | DorsAttn | Rest vs 2-Back | 3.46 | 0.0096 | 32 |
| 73 | DorsAttn | Rest vs 0-Back | -3.50 | 0.0050 | 34 |
| 73 | DorsAttn | Rest vs 1-Back | -3.42 | 0.0050 | 34 |
| 75 | SalVentAttn | Rest vs 2-Back | 3.32 | 0.0139 | 32 |
| 75 | SalVentAttn | 0-Back vs 2-Back | 2.38 | 0.0477 | 32 |
| 75 | SalVentAttn | 1-Back vs 2-Back | 2.60 | 0.0430 | 32 |
| 78 | SalVentAttn | Rest vs 1-Back | -2.68 | 0.0351 | 32 |
| 78 | SalVentAttn | 1-Back vs 2-Back | 2.69 | 0.0351 | 32 |
| 79 | Limbic | Rest vs 0-Back | -5.60 | 1.16e-05 | 32 |
| 79 | Limbic | Rest vs 1-Back | -6.39 | 2.47e-06 | 32 |
| 79 | Limbic | Rest vs 2-Back | -4.91 | 5.61e-05 | 32 |
| 80 | Limbic | Rest vs 0-Back | -2.40 | 0.0440 | 34 |
| 80 | Limbic | Rest vs 1-Back | -3.97 | 0.0022 | 34 |
| 80 | Limbic | 1-Back vs 2-Back | 2.98 | 0.0160 | 34 |
| 83 | Cont | Rest vs 0-Back | -3.57 | 0.0027 | 34 |

| | | | | | |
|---|---|---|---|---|---|
| 83 | Cont | Rest vs 1-Back | -4.93 | 1.35e-04 | 34 |
| 83 | Cont | Rest vs 2-Back | -3.51 | 0.0027 | 34 |
| 86 | Cont | Rest vs 0-Back | -2.84 | 0.0229 | 34 |
| 86 | Cont | Rest vs 1-Back | -3.45 | 0.0093 | 34 |
| 86 | Cont | Rest vs 2-Back | -2.51 | 0.0344 | 34 |
| 87 | Cont | Rest vs 0-Back | -2.55 | 0.0325 | 30 |
| 87 | Cont | Rest vs 1-Back | -2.88 | 0.0220 | 30 |
| 87 | Cont | 1-Back vs 2-Back | 3.05 | 0.0220 | 30 |
| 88 | Cont | Rest vs 0-Back | -3.15 | 0.0070 | 34 |
| 88 | Cont | Rest vs 1-Back | -3.85 | 0.0031 | 34 |
| 88 | Cont | Rest vs 2-Back | -3.23 | 0.0070 | 34 |
| 91 | Default | Rest vs 0-Back | -3.71 | 0.0025 | 33 |
| 91 | Default | Rest vs 1-Back | -3.70 | 0.0025 | 33 |
| 92 | Default | Rest vs 1-Back | -2.57 | 0.0443 | 34 |
| 92 | Default | 1-Back vs 2-Back | 2.71 | 0.0443 | 34 |
| 94 | Default | Rest vs 0-Back | -5.30 | 2.26e-05 | 34 |
| 94 | Default | Rest vs 1-Back | -6.21 | 3.15e-06 | 34 |
| 94 | Default | Rest vs 2-Back | -3.98 | 7.14e-04 | 34 |
| 96 | Default | 0-Back vs 2-Back | 3.06 | 0.0133 | 33 |
| 96 | Default | 1-Back vs 2-Back | 3.25 | 0.0133 | 33 |
| 97 | Default | Rest vs 0-Back | -3.83 | 0.0017 | 33 |
| 97 | Default | Rest vs 1-Back | -4.00 | 0.0017 | 33 |
| 97 | Default | Rest vs 2-Back | -2.65 | 0.0248 | 33 |
| 97 | Default | 1-Back vs 2-Back | 2.51 | 0.0257 | 33 |
| 99 | Default | Rest vs 2-Back | 3.76 | 0.0044 | 31 |
| 99 | Default | 0-Back vs 2-Back | 3.36 | 0.0064 | 31 |
| 99 | Default | 1-Back vs 2-Back | 2.71 | 0.0221 | 31 |
| 100 | Default | Rest vs 1-Back | 2.23 | 0.0493 | 33 |
| 100 | Default | Rest vs 2-Back | 5.06 | 9.93e-05 | 33 |
| 100 | Default | 0-Back vs 2-Back | 3.80 | 0.0018 | 33 |
| 100 | Default | 1-Back vs 2-Back | 3.53 | 0.0026 | 33 |

**Table S2.** Parcel-level post-hoc pairwise t-tests between sessions, FDR-corrected; only significant comparisons shown.

#### Within-network tests across sessions

A two-way repeated-measures ANOVA (Network × Session) confirmed that cognitive state systematically modulated network-level differences, with robust main effects of Network ($F(6,198)=77.01$, $p_{GG\text{-}corr}<0.001$, partial $\eta^2=0.70$) and Session ($F(3,99)=3.11$, $p_{GG\text{-}corr}=0.04$, partial $\eta^2=0.09$), and a significant Network × Session interaction ($F(18,594)=13.15$, $p_{GG\text{-}corr}<0.001$, partial $\eta^2=0.28$), indicating that the effect of cognitive state on cortical persistence differed across networks.

Network-averaged effect sizes of the parcel-wise ANOVA further highlighted LIM and DMN as the most sensitive to task-driven H modulation (LIM=0.26±0.12; DMN=0.17±0.09). Pairwise between-network contrasts (Table S5, S6) indicate that state-dependent reconfiguration of cortical persistence is not uniform but concentrated within LIM and DMN, the networks showing the largest rest-to-task H shifts, with task engagement reordering networks' relative standing rather than uniformly raising or lowering persistence (Supplementary Table S7, S8).

To determine which networks showed significant session-wise modulation of H, we ran a Friedman test across the four sessions (Rest, 0-Back, 1-Back, 2-Back) for each of the seven Yeo networks, followed by post-hoc pairwise Wilcoxon signed-rank tests for networks with a significant omnibus effect. Both omnibus and pairwise p-values are FDR-corrected across networks and, for pairwise tests, across session pairs.

| Network | n | Friedman $\chi^2$ | p (raw) | p (FDR) |
|---|---|---|---|---|
| Vis | 34 | 5.65 | 0.130 | 0.229 |
| SomMot | 34 | 5.64 | 0.131 | 0.229 |
| DorsAttn | 34 | 1.88 | 0.597 | 0.597 |
| SalVentAttn | 34 | 4.08 | 0.253 | 0.354 |
| Limbic | 34 | 40.91 | 6.83e-09 | 4.78e-08 |
| Cont | 34 | 2.24 | 0.524 | 0.597 |
| Default | 34 | 13.09 | 0.004 | 0.016 |

**Table S3.** Omnibus Friedman tests of H across sessions, within each Yeo-Krienen network.

| Network | Comparison | n | Wilcoxon W | p (raw) | p (FDR) |
|---|---|---|---|---|---|
| Limbic | Rest vs 0-Back | 35 | 38.0 | 3.839e-07 | 2.304e-06 |
| Limbic | Rest vs 1-Back | 35 | 23.0 | 3.725e-08 | 4.470e-07 |
| Limbic | Rest vs 2-Back | 34 | 105.0 | 6.278e-04 | 0.003 |
| Limbic | 0-Back vs 1-Back | 35 | 257.0 | 0.489 | 0.556 |
| Limbic | 0-Back vs 2-Back | 34 | 176.0 | 0.037 | 0.064 |
| Limbic | 1-Back vs 2-Back | 34 | 136.0 | 0.005 | 0.010 |
| Default | Rest vs 0-Back | 35 | 220.0 | 0.123 | 0.169 |
| Default | Rest vs 1-Back | 35 | 221.0 | 0.127 | 0.169 |
| Default | Rest vs 2-Back | 34 | 258.0 | 0.510 | 0.556 |
| Default | 0-Back vs 1-Back | 35 | 288.0 | 0.871 | 0.871 |
| Default | 0-Back vs 2-Back | 34 | 134.0 | 0.004 | 0.010 |
| Default | 1-Back vs 2-Back | 34 | 113.0 | 0.001 | 0.003 |

**Table S4.** Post-hoc pairwise Wilcoxon signed-rank tests between sessions, within each network with a significant omnibus effect.

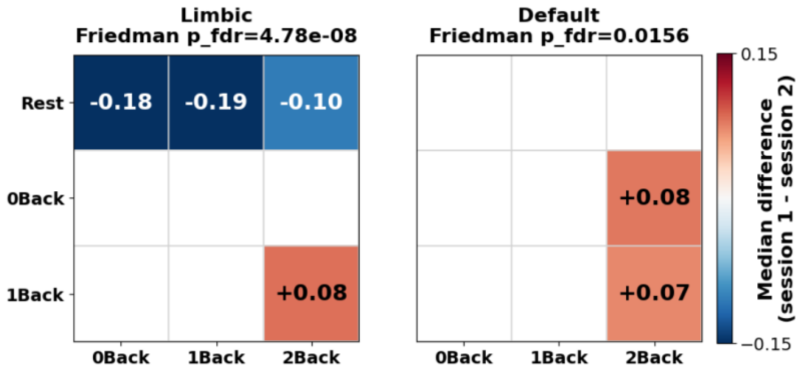


**Figure S7.** Session-wise median H differences for the two networks (Limbic, Default Mode) showing a significant omnibus Friedman effect.

#### Within-session tests across networks

As a complementary, session-anchored view of the same effect, we next tested whether H differed across the seven Yeo networks within each session (omnibus Friedman test), followed by post-hoc pairwise Wilcoxon tests between all network pairs; only pairs surviving FDR correction are listed in the table below.

| Session | n | Friedman $\chi^2$ | p (raw) | p (FDR) |
|---|---|---|---|---|
| Rest | 35 | 127.151 | 5.113e-25 | 2.045e-24 |
| 0Back | 35 | 107.363 | 7.264e-21 | 7.264e-21 |
| 1Back | 35 | 108.086 | 5.129e-21 | 6.838e-21 |
| 2Back | 34 | 118.727 | 3.016e-23 | 6.032e-23 |

**Table S5.** Omnibus Friedman tests of H across the seven Yeo-Krienen networks, within each session.

| Session | Comparison | n | Wilcoxon W | p (raw) | p (FDR) |
|---|---|---|---|---|---|
| Rest | Vis vs SomMot | 35 | 173.000 | 0.019 | 0.024 |
| Rest | Vis vs DorsAttn | 35 | 10.000 | 2.503e-09 | 1.051e-08 |
| Rest | Vis vs Limbic | 35 | 5.000 | 5.821e-10 | 4.445e-09 |
| Rest | Vis vs Cont | 35 | 31.000 | 1.382e-07 | 3.629e-07 |
| Rest | Vis vs Default | 35 | 94.000 | 1.392e-04 | 2.339e-04 |
| Rest | SomMot vs DorsAttn | 35 | 60.000 | 5.685e-06 | 1.137e-05 |
| Rest | SomMot vs Limbic | 35 | 0.000 | 5.821e-11 | 9.779e-10 |
| Rest | SomMot vs Cont | 35 | 130.000 | 0.002 | 0.003 |
| Rest | DorsAttn vs SalVentAttn | 35 | 19.000 | 1.787e-08 | 6.004e-08 |
| Rest | DorsAttn vs Limbic | 35 | 0.000 | 5.821e-11 | 9.779e-10 |
| Rest | DorsAttn vs Cont | 35 | 149.000 | 0.006 | 0.008 |
| Rest | DorsAttn vs Default | 35 | 52.000 | 2.298e-06 | 4.948e-06 |
| Rest | SalVentAttn vs Limbic | 35 | 1.000 | 1.164e-10 | 1.087e-09 |
| Rest | SalVentAttn vs Cont | 35 | 32.000 | 1.609e-07 | 4.097e-07 |
| Rest | SalVentAttn vs Default | 35 | 131.000 | 0.002 | 0.003 |
| Rest | Limbic vs Cont | 35 | 0.000 | 5.821e-11 | 9.779e-10 |
| Rest | Limbic vs Default | 35 | 1.000 | 1.164e-10 | 1.087e-09 |
| Rest | Cont vs Default | 35 | 62.000 | 7.053e-06 | 1.346e-05 |
| 0Back | Vis vs DorsAttn | 35 | 0.000 | 5.821e-11 | 9.779e-10 |
| 0Back | Vis vs Cont | 35 | 1.000 | 1.164e-10 | 1.087e-09 |
| 0Back | Vis vs Default | 35 | 11.000 | 3.201e-09 | 1.222e-08 |
| 0Back | SomMot vs DorsAttn | 35 | 31.000 | 1.382e-07 | 3.629e-07 |
| 0Back | SomMot vs Limbic | 35 | 141.000 | 0.004 | 0.005 |
| 0Back | SomMot vs Cont | 35 | 60.000 | 5.685e-06 | 1.137e-05 |
| 0Back | SomMot vs Default | 35 | 107.000 | 3.828e-04 | 6.067e-04 |
| 0Back | DorsAttn vs SalVentAttn | 35 | 9.000 | 1.921e-09 | 8.964e-09 |
| 0Back | DorsAttn vs Limbic | 35 | 30.000 | 1.185e-07 | 3.317e-07 |
| 0Back | SalVentAttn vs Limbic | 35 | 144.000 | 0.004 | 0.006 |
| 0Back | SalVentAttn vs Cont | 35 | 35.000 | 2.509e-07 | 5.854e-07 |

| | | | | | |
|---|---|---|---|---|---|
| 0Back | SalVentAttn vs Default | 35 | 88.000 | 8.434e-05 | 1.476e-04 |
| 0Back | Limbic vs Cont | 35 | 9.000 | 1.921e-09 | 8.964e-09 |
| 0Back | Limbic vs Default | 35 | 11.000 | 3.201e-09 | 1.222e-08 |
| 0Back | Cont vs Default | 35 | 154.000 | 0.007 | 0.010 |
| 1Back | Vis vs SomMot | 35 | 126.000 | 0.001 | 0.002 |
| 1Back | Vis vs DorsAttn | 35 | 6.000 | 8.149e-10 | 4.889e-09 |
| 1Back | Vis vs SalVentAttn | 35 | 162.000 | 0.011 | 0.014 |
| 1Back | Vis vs Cont | 35 | 0.000 | 5.821e-11 | 9.779e-10 |
| 1Back | Vis vs Default | 35 | 7.000 | 1.106e-09 | 6.112e-09 |
| 1Back | SomMot vs DorsAttn | 35 | 66.000 | 1.073e-05 | 2.002e-05 |
| 1Back | SomMot vs Limbic | 35 | 120.000 | 9.603e-04 | 0.001 |
| 1Back | SomMot vs Cont | 35 | 34.000 | 2.168e-07 | 5.204e-07 |
| 1Back | SomMot vs Default | 35 | 126.000 | 0.001 | 0.002 |
| 1Back | DorsAttn vs SalVentAttn | 35 | 20.000 | 2.160e-08 | 6.718e-08 |
| 1Back | DorsAttn vs Limbic | 35 | 44.000 | 8.570e-07 | 1.894e-06 |
| 1Back | DorsAttn vs Cont | 35 | 103.000 | 2.834e-04 | 4.577e-04 |
| 1Back | SalVentAttn vs Limbic | 35 | 117.000 | 7.825e-04 | 0.001 |
| 1Back | SalVentAttn vs Cont | 35 | 10.000 | 2.503e-09 | 1.051e-08 |
| 1Back | SalVentAttn vs Default | 35 | 61.000 | 6.335e-06 | 1.238e-05 |
| 1Back | Limbic vs Cont | 35 | 20.000 | 2.160e-08 | 6.718e-08 |
| 1Back | Limbic vs Default | 35 | 41.000 | 5.779e-07 | 1.312e-06 |
| 1Back | Cont vs Default | 35 | 60.000 | 5.685e-06 | 1.137e-05 |
| 2Back | Vis vs DorsAttn | 34 | 4.000 | 8.149e-10 | 4.889e-09 |
| 2Back | Vis vs Cont | 34 | 11.000 | 6.403e-09 | 2.338e-08 |
| 2Back | Vis vs Default | 34 | 73.000 | 4.236e-05 | 7.570e-05 |
| 2Back | SomMot vs DorsAttn | 34 | 25.000 | 1.052e-07 | 3.048e-07 |
| 2Back | SomMot vs Limbic | 34 | 93.000 | 2.475e-04 | 4.077e-04 |
| 2Back | SomMot vs Cont | 34 | 14.000 | 1.281e-08 | 4.482e-08 |
| 2Back | SomMot vs Default | 34 | 157.000 | 0.015 | 0.019 |
| 2Back | DorsAttn vs SalVentAttn | 34 | 17.000 | 2.410e-08 | 7.229e-08 |
| 2Back | DorsAttn vs Limbic | 34 | 5.000 | 1.164e-09 | 6.112e-09 |
| 2Back | DorsAttn vs Cont | 34 | 156.000 | 0.015 | 0.019 |
| 2Back | DorsAttn vs Default | 34 | 64.000 | 1.723e-05 | 3.146e-05 |
| 2Back | SalVentAttn vs Limbic | 34 | 130.000 | 0.003 | 0.005 |
| 2Back | SalVentAttn vs Cont | 34 | 4.000 | 8.149e-10 | 4.889e-09 |
| 2Back | SalVentAttn vs Default | 34 | 81.000 | 8.892e-05 | 1.524e-04 |
| 2Back | Limbic vs Cont | 34 | 0.000 | 1.164e-10 | 1.087e-09 |
| 2Back | Limbic vs Default | 34 | 29.000 | 2.024e-07 | 5.002e-07 |

**Table S6.** Post-hoc pairwise Wilcoxon signed-rank tests between networks, within each session; only FDR-significant comparisons shown.

##### Tests on ΔH matrices across sessions

Finally, to test whether the magnitude of between-network differences in H changed across cognitive states, we computed the session-wise ΔH (the difference in mean H between each pair of networks) and ran a Friedman test across the four sessions on this ΔH series; network pairs with a significant omnibus effect were followed up with pairwise Wilcoxon tests between sessions, again FDR-corrected and reported below only where significant.

| Network 1 | Network 2 | n | Friedman χ² | P (raw) | p (FDR) |
|---|---|---|---|---|---|
| Vis | SomMot | 34 | 4.221 | 0.239 | 0.278 |
| Vis | DorsAttn | 34 | 1.195 | 0.754 | 0.754 |
| Vis | SalVentAttn | 34 | 2.628 | 0.453 | 0.500 |
| Vis | Limbic | 34 | 33.690 | 2.30e-07 | 1.21e-06 |
| Vis | Cont | 34 | 16.646 | 8.36e-04 | 0.002 |
| Vis | Default | 34 | 6.611 | 0.085 | 0.120 |
| SomMot | DorsAttn | 34 | 4.681 | 0.197 | 0.243 |
| SomMot | SalVentAttn | 34 | 1.407 | 0.704 | 0.739 |
| SomMot | Limbic | 34 | 47.336 | 2.95e-10 | 6.19e-09 |
| SomMot | Cont | 34 | 18.239 | 3.93e-04 | 0.001 |
| SomMot | Default | 34 | 6.239 | 0.101 | 0.132 |
| DorsAttn | SalVentAttn | 34 | 7.088 | 0.069 | 0.104 |
| DorsAttn | Limbic | 34 | 43.425 | 2.00e-09 | 2.10e-08 |
| DorsAttn | Cont | 34 | 16.416 | 9.32e-04 | 0.002 |
| DorsAttn | Default | 34 | 16.363 | 9.55e-04 | 0.002 |
| SalVentAttn | Limbic | 34 | 34.345 | 1.68e-07 | 1.17e-06 |
| SalVentAttn | Cont | 34 | 17.425 | 5.78e-04 | 0.001 |
| SalVentAttn | Default | 34 | 11.513 | 0.009 | 0.015 |
| Limbic | Cont | 34 | 31.124 | 8.00e-07 | 3.36e-06 |
| Limbic | Default | 34 | 26.257 | 8.43e-06 | 2.53e-05 |
| Cont | Default | 34 | 28.628 | 2.68e-06 | 9.38e-06 |

**Table S7.** Omnibus Friedman tests of the session-wise ΔH (between-network H difference) across sessions, for each network pair.

| Network 1 | Network 2 | Session comparison | n | Wilcoxon W | p (raw) | p (FDR) |
|---|---|---|---|---|---|---|
| Vis | Limbic | Rest vs 0Back | 35 | 38 | 3.84e-07 | 2.72e-06 |
| Vis | Limbic | Rest vs 1Back | 35 | 35 | 2.51e-07 | 2.45e-06 |
| Vis | Limbic | Rest vs 2Back | 34 | 42 | 1.32e-06 | 7.34e-06 |
| Vis | Cont | Rest vs 1Back | 35 | 156 | 0.008 | 0.019 |
| Vis | Cont | Rest vs 2Back | 34 | 146 | 0.009 | 0.019 |
| Vis | Cont | 0Back vs 1Back | 35 | 139 | 0.007 | 0.016 |
| Vis | Cont | 0Back vs 2Back | 34 | 137 | 0.005 | 0.013 |
| SomMot | Limbic | Rest vs 0Back | 35 | 4 | 4.07e-10 | 3.18e-08 |
| SomMot | Limbic | Rest vs 1Back | 35 | 27 | 7.34e-08 | 1.15e-06 |
| SomMot | Limbic | Rest vs 2Back | 34 | 33 | 3.74e-07 | 2.72e-06 |
| SomMot | Cont | Rest vs 1Back | 35 | 129 | 0.002 | 0.005 |
| SomMot | Cont | Rest vs 2Back | 34 | 98 | 3.69e-04 | 0.001 |
| DorsAttn | Limbic | Rest vs 0Back | 35 | 8 | 1.46e-09 | 5.68e-08 |
| DorsAttn | Limbic | Rest vs 1Back | 35 | 19 | 1.79e-08 | 3.48e-07 |
| DorsAttn | Limbic | Rest vs 2Back | 34 | 58 | 9.05e-06 | 3.92e-05 |
| DorsAttn | Cont | Rest vs 1Back | 35 | 87 | 7.74e-05 | 3.02e-04 |
| DorsAttn | Cont | Rest vs 2Back | 34 | 96 | 3.15e-04 | 0.001 |
| DorsAttn | Cont | 0Back vs 1Back | 35 | 137 | 0.006 | 0.015 |

| | | | | | | |
|---|---|---|---|---|---|---|
| DorsAttn | Default | Rest vs 0Back | 35 | 157 | 0.009 | 0.019 |
| DorsAttn | Default | Rest vs 1Back | 35 | 129 | 0.002 | 0.005 |
| DorsAttn | Default | 0Back vs 2Back | 34 | 157 | 0.015 | 0.033 |
| DorsAttn | Default | 1Back vs 2Back | 34 | 84 | 1.16e-04 | 4.11e-04 |
| SalVentAttn | Limbic | Rest vs 0Back | 35 | 30 | 1.18e-07 | 1.54e-06 |
| SalVentAttn | Limbic | Rest vs 1Back | 35 | 35 | 2.51e-07 | 2.45e-06 |
| SalVentAttn | Limbic | Rest vs 2Back | 34 | 55 | 6.47e-06 | 3.15e-05 |
| SalVentAttn | Cont | Rest vs 2Back | 34 | 109 | 8.40e-04 | 0.003 |
| SalVentAttn | Cont | 0Back vs 1Back | 35 | 164 | 0.022 | 0.047 |
| SalVentAttn | Cont | 0Back vs 2Back | 34 | 112 | 0.001 | 0.003 |
| Limbic | Cont | Rest vs 0Back | 35 | 17 | 1.20e-08 | 3.13e-07 |
| Limbic | Cont | Rest vs 1Back | 35 | 63 | 7.84e-06 | 3.60e-05 |
| Limbic | Cont | Rest vs 2Back | 34 | 117 | 0.001 | 0.004 |
| Limbic | Default | Rest vs 0Back | 35 | 40 | 5.05e-07 | 3.28e-06 |
| Limbic | Default | Rest vs 1Back | 35 | 46 | 1.11e-06 | 6.64e-06 |
| Limbic | Default | Rest vs 2Back | 34 | 44 | 1.71e-06 | 8.89e-06 |
| Cont | Default | Rest vs 2Back | 34 | 81 | 8.89e-05 | 3.30e-04 |
| Cont | Default | 0Back vs 2Back | 34 | 32 | 3.22e-07 | 2.72e-06 |
| Cont | Default | 1Back vs 2Back | 34 | 75 | 5.12e-05 | 2.10e-04 |

**Table S8.** Post-hoc pairwise Wilcoxon signed-rank tests on ΔH between sessions, for each network pair; only FDR-significant comparisons shown.

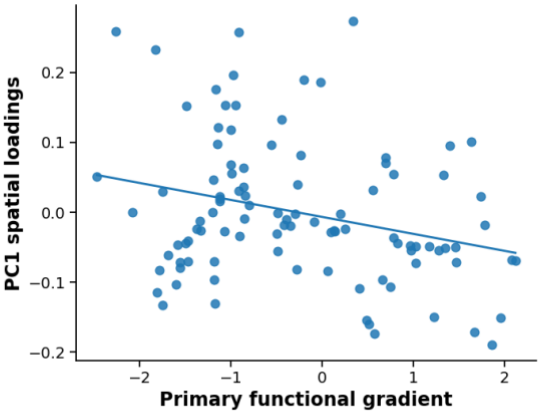


**Figure S8.** Scatterplot and regression line showing the parcel-wise spatial correspondence between PC1 loadings and the primary functional gradient.

#### Alternative parcellation (Glasser360) replication

The mean parcel-wise Hurst maps and network-averaged radar plots in both the discovery (Figure S8) and replication (Figure S9A-B) datasets showed spatial patterns qualitatively consistent with those obtained using the Schaefer parcellation. H values were slightly lower overall, potentially reflecting noisier signals associated with the smaller parcel size. Mean H remained comparable across datasets and showed the same non-monotonic modulation across cognitive load in the replication sample (discovery: Rest = 1.20 ± 0.24; replication: Rest = 1.17 ± 0.16, 0-Back = 1.22 ± 0.14, 1-Back = 1.22 ± 0.14, 2-Back = 1.15 ± 0.14). The Limbic network (LIM) showed the lowest H at rest in both datasets (discovery: H = 0.98 ± 0.08; replication: H = 0.92 ± 0.21), whereas the Control (CONT) and Default Mode (DMN) networks remained among the networks with the highest H values (discovery: CONT = 1.27 ± 0.14,

DMN = 1.24 ± 0.13; replication, mean across sessions: CONT = 1.29 ± 0.25, DMN = 1.24 ± 0.25).

In the replication dataset, parcel-wise ANOVA revealed a significant effect of session in 66% of parcels, with the strongest effects observed in LIM and DMN (Figure S9C). The SVD analysis also reproduced the main patterns observed with the primary parcellation (Figure S9D). The first component, explaining 47% of the variance, showed positive loadings across all sessions and a spatial profile characterized by positive loadings in CONT, DMN, and DAN and negative loadings in LIM. The second component, explaining 6% of the variance, captured an increase from rest to task conditions and showed positive loadings in LIM and CONT and negative loadings in VIS.

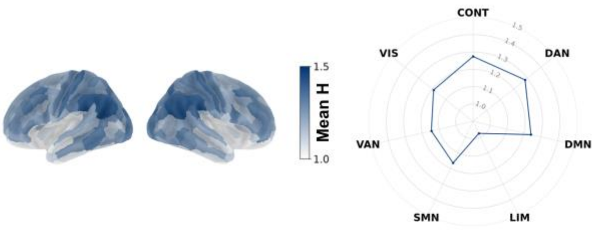


**Figure S9.** Mean parcel-wise Hurst exponent maps at rest in the discovery dataset and corresponding network-averaged mean H using Glasser360 parcellation.

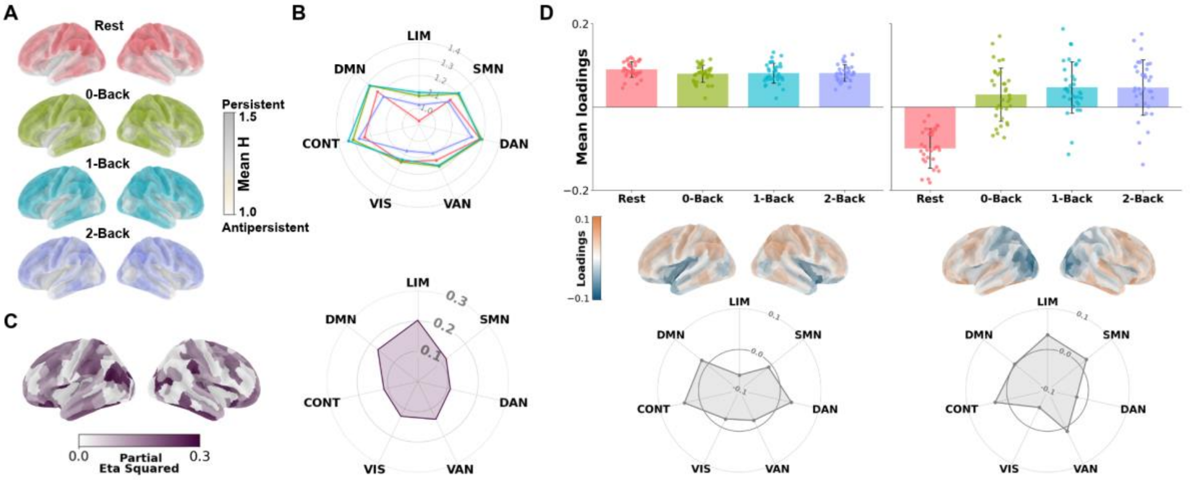


**Figure S10. Alternative parcellation (Glasser360) replication of Figure 3. A.** Mean parcel-wise Hurst exponent maps across the four sessions. **B.** Network-averaged mean H displayed as radar plots for the seven functional networks across the four sessions. Shaded areas in the radar plot indicate 95% confidence intervals. **C.** Parcel-wise partial eta squared maps from the repeated-measures ANOVA, together with the corresponding network-level summary of effect sizes. **D.** Singular value decomposition of the parcel × (subject·session) H matrix, showing the mean session loadings of the first two dominant components, their associated spatial loading maps and the mean network-wise loadings as radar plots. Abbreviations: VIS, Visual; SMN, Somatomotor; DAN, Dorsal Attention; VAN, Salience/Ventral Attention; LIM, Limbic; CONT, Control; DMN, Default Mode Network.

Cytoarchitectural voxel composition

| Voxel | Parcel | Von Economo class | BigBrain gradient |
|---|---|---|---|
| Anterior Cingulate | 28 | 1 | 0.0373 |
| | 78 | 1 | 0.0351 |
| | 88 | 1 | 0.0396 |
| Occipital | 6 | 5 | -0.0310 |
| | 9 | 3 | -0.0680 |
| | 36 | 2 | -0.0031 |
| | 49 | 2 | 0.0252 |
| | 50 | 2 | 0.0234 |
| | 55 | 5 | -0.0659 |
| | 99 | 3 | -0.0051 |
| | 100 | 2 | -0.0065 |
| Right Auditory | 59 | 3 | -0.0085 |
| | 92 | 3 | 0.0077 |
| | 93 | 2 | 0.0355 |
| Somatomotor Left | 14 | 1 | -0.0877 |
| | 19 | 2 | 0.0554 |

**Table S9.** Cytoarchitectural composition of each MRS voxel: constituent Schaefer parcels, von Economo class, and BigBrain gradient value.

#### Task-dependent reconfiguration of the H–functional gradient relationship

Both linear and quadratic gradient components differed significantly from rest at every load (0-Back: linear β=0.009, t=7.94, p<0.001; quadratic β=−0.0013, t=−3.01, p=0.003; 1-Back: linear β=0.011, t=9.42, p<0.001; quadratic β=−0.0012, t=−2.73, p=0.006; 2-Back: linear β=0.009, t=7.44, p<0.001; quadratic β=−0.0022, t=−5.06, p<0.001), indicating systematic reconfiguration of the gradient–H relationship with load. Simple-slope tests showed the quadratic component was significant at rest, 0-Back, and 1-Back but not 2-Back, whereas the linear component was significant at 0-Back, 1-Back, and 2-Back but not rest.

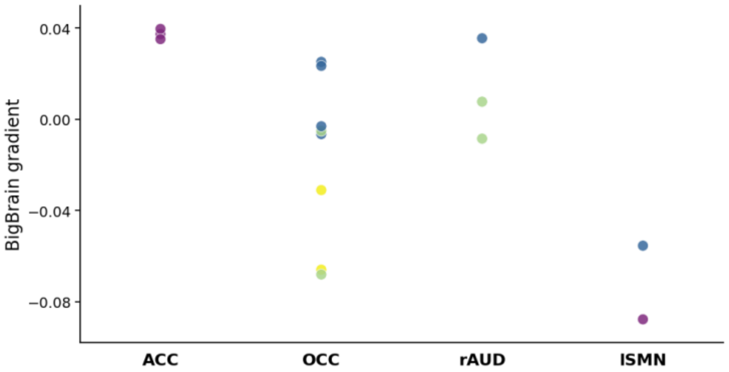


**Figure S11.** BigBrain gradient values of the parcels composing each MRS voxel (ACC, OCC, rAUD, ISMN), coloured by von Economo class.

#### Test-retest reliability of H in HCP

H maps were highly reproducible across sessions (group maps, REST1 vs REST2: spatial r=0.99; single runs: 0.93-0.98) and moderately reliable for individual differences (per-parcel ICC(3,1), median 0.59, IQR 0.55-0.62; subject mean H, REST1 vs REST2: r=0.61). Individual topographies were stable across days (median spatial r=0.81) and individually distinctive: deviations from the group map correlated at a median r=0.63 for the same subject across days versus 0.00 across different subjects, and a participant's session-2 map could be identified from their session-1 map in 64% of cases (chance 1%).
